# Explaining distortions in metacognition with an attractor network model of decision uncertainty

**DOI:** 10.1101/2020.09.25.313619

**Authors:** Nadim A. A. Atiya, Quentin J. M. Huys, Raymond J. Dolan, Stephen M. Fleming

## Abstract

Metacognition is the ability to reflect on, and evaluate, our cognition and behaviour. Distortions in metacognition are common in mental health disorders, though the neural underpinnings of such dysfunction are unknown. One reason for this is that models of key components of metacognition, such as decision confidence, are generally specified at an algorithmic or process level. While such models can be used to relate brain function to psychopathology, they are difficult to map to a neurobiological mechanism. Here, we develop a biologically-plausible model of decision uncertainty in an attempt to bridge this gap. We first relate the model’s uncertainty in perceptual decisions to standard metrics of metacognition, namely mean confidence level (bias) and the accuracy of metacognitive judgments (sensitivity). We show that dissociable shifts in metacognition are associated with isolated disturbances at higher-order levels of a circuit associated with self-monitoring, akin to neuropsychological findings that highlight the detrimental effect of prefrontal brain lesions on metacognitive performance. Notably, we are able to account for empirical confidence judgements by fitting the parameters of our biophysical model to first-order performance data, specifically choice and response times. Lastly, in a reanalysis of existing data we show that self-reported mental health symptoms relate to disturbances in an uncertainty-monitoring component of the network. By bridging a gap between a biologically-plausible model of confidence formation and observed disturbances of metacognition in mental health disorders we provide a first step towards mapping theoretical constructs of metacognition onto dynamical models of decision uncertainty. In doing so, we provide a computational framework for modelling metacognitive performance in settings where access to explicit confidence reports is not possible.

**Author Summary:** In this work, we use a biologically-plausible model of decision uncertainty to show that shifts in metacognition are associated with disturbances in the interaction between decision-making and higher-order uncertainty-monitoring networks. Specifically, we show that stronger uncertainty modulation is associated with decreased metacognitive bias, sensitivity, and efficiency, with no effect on perceptual sensitivity. Our approach not only enables inferences about uncertainty modulation (and, in turn, these facets of metacognition) from fits to first-order performance data alone – but also provides a first step towards relating dynamical models of decision-making to metacognition. We also relate our model’s uncertainty modulation to psychopathology, and show that it can offer an implicit, low-dimensional marker of metacognitive (dys)function – opening the door to richer analysis of the interaction between metacognitive performance and psychopathology from first-order performance data.

## Introduction

Computational psychiatry (Friston et al. 2014; Huys, Maia, and Frank 2016; Wang and Krystal 2014; Montague et al. 2012) employs mechanistic and theory-driven models to relate brain function to phenomena that characterise mental health disorders (Ratcliff 1978; Ratcliff, Smith, and McKoon 2015; Rescorla, Wagner et al. 1972; Huys, Maia, and Frank 2016; Sutton and Barto 2018). Typically, algorithmic-level models (Marr and Poggio 1976) describe the computational processes that realise specific brain functions and return theoretically meaningful parameters that may vary between subjects. Some of these algorithmic models (e.g. reinforcement learning; Sutton and Barto 2018) closely relate to the functions of discrete brain circuits (Schultz 1999; Dayan and Balleine 2002; Dolan and Dayan 2012). However, there remains a high degree of imprecision when relating diverse sets of algorithms to circuit-level disturbances, potentially limiting our understanding of, and treatments for, mental disorders.

One proposal is that the same neural circuit disturbances can be associated with several (often unrelated) changes in behaviour (Stephan et al. 2016). Here detailed biophysical models (Murray et al. 2014; Krystal et al. 2017; Rolls, Loh, and Deco 2008) may provide tools for understanding mental health disorders in terms of precise disturbances at the microcircuit level. For instance, Murray et al. (2014) showed that an imbalance in excitatory/inhibitory synaptic connections in a spiking neural network model can explain working memory deficits associated with schizophrenia. However, the complex nature of such models renders it challenging to fit them to individual subjects’ behavioural data. At the level of neural systems, simpler biologically-grounded models (Dima et al. 2009; Yang et al. 2014) have been employed to relate macrocircuit-level dysfunctions to symptoms of mental health disorders, and motivate non-invasive experimental neuroimaging to probe such dysfunctions (Cohen and Servan-Schreiber 1992). Such (connectionist) biologically-motivated models retain a mapping between neurobiology and behaviour, while allowing faster computation and fewer free parameters.

Here our focus is on developing similar biologically-plausible models of subjective confidence and metacognition – the ability to reflect upon and evaluate aspects of our own cognition and behaviour. Recent advances in metacognition research has led to the development of precision assays for different facets of metacognitive ability (Maniscalco and Lau 2012; Fleming 2017). Within a signal detection theory (SDT) framework, metacognitive bias refers to a subject’s overall (mean) confidence level on a task. In contrast, metacognitive sensitivity refers to whether subjects’ confidence ratings effectively distinguish between correct and incorrect decisions, as quantified by the SDT metric *meta – d′*. Furthermore, metacognitive sensitivity can be compared to another SDT measure, d′, which quantifies how effectively a subject processes information related to the task (Howell 2009; Rounis et al. 2010). The ratio *meta* – *d′/d′* thus yields a measure of metacognitive efficiency, i.e. metacognitive sensitivity for a given level of task performance (Fleming and Lau 2014).

Experimental evidence suggests that these facets of metacognitive ability are dissociable from task performance, and may have a distinct neural and computational basis (Del Cul et al. 2009; Fleming et al. 2010; Fleming and Dolan 2012; Fleming et al., 2014; Lak et al., 2014; Bang & Fleming, 2018; Miyamoto et al., 2018). Interestingly, self-reported mental health symptoms have been linked to changes in metacognition, often in the absence of differences in task performance (Rouault et al. 2018; Moses-Payne et al. 2019; Hoven et al., 2019; Seow & Gillan, 2020). Developing a biologically-motivated model of metacognition has the potential to cast light on how this dissociable mechanism is implemented at a circuit level, as well as provide a direct bridge between circuit-level dysfunction and psychopathology.

Theoretical work addressing perceptual decision-making has proposed dynamical reduced accounts (Wong and Wang 2006; Roxin and Ledberg 2008) that provide detailed biophysical models of decision making (Wang 2002), enabling more rigorous theoretical analyses and faster computation. For instance, Wong and Wang (2006) have accounted for most of the behavioural results addressed by Wang (2002)’s model using the two slowest N-Methyl-D-aspartic acid (NMDA) dynamical variables. More recently, Atiya et al. (2019) extended Wong and Wang (2006)’s model to account for decision confidence reports and other metacognitive behaviours, such as an ability to flexibly change one’s mind and correct errors (Atiya et al. 2020). More specifically, guided by neurophysiological evidence that supports an encoding of confidence within higher-order prefrontal brain regions (Kepecs et al. 2008; Fleming and Dolan 2012), the authors introduced the idea of a third ‘uncertainty-monitoring’ neuronal population (i.e. dynamical variable). This population continuously monitors uncertainty in the network, interacting with the other two populations involved in decision-making via a feedback loop mechanism (Yeung et al., 2004).

A classic proposal from cognitive psychology is that changes in metacognition reflect alterations in higher-order computations that serve to “monitor” first-order task performance (Nelson & Narens, 1990). Our primary focus here is on the question of whether developing biologically-plausible accounts of metacognitive monitoring shed light on the source of differences metacognitive sensitivity. Other recent work has focused on simulating parallel neural populations engaged in perceptual decision-making, finding that informing confidence with the activity of less-normalisation-tuned neurons can account for cases in which confidence is altered in the absence of differences in performance (Maniscalco et al., 2021, a shift in metacognitive bias). Our model is complementary to this endeavour, instead focusing on the dynamics of uncertainty encoding within a dedicated, higher-order neural population that integrates input from sensorimotor neuronal pools, and continuously feeds this uncertainty signal back to modulate evidence integration. This feedback mechanism adds a layer of nonlinearity, accounting for non-trivial interactions between confidence, accuracy and response times. We will see, though, that such a higher-order monitoring population can also account for shifts in metacognitive bias, and therefore capture instances of performance-confidence dissociation.

To gain insight into potential mechanisms underlying shifts in metacognition, we first demonstrate that our biologically-motivated model (Atiya et al. 2019, 2020) can account for human confidence reports. In a novel approach, we show that the intrinsic dynamics of this model, constrained only by first-order performance, are sufficient to account for subjects’ confidence reports, going beyond existing methods of fitting models directly to empirical confidence data (Pleskac and Busemeyer 2010; Sanders, Hangya, and Kepecs 2016). We then map theoretical constructs such as metacognitive sensitivity and efficiency onto our dynamical model, demonstrating that changes in metacognitive sensitivity are associated with isolated disturbances in a higher-order node of the network involved in uncertainty monitoring. This computational approach also allowed us to relate circuit-level deficits in metacognition to psychopathology, by re-analysing an existing dataset (Rouault et al., 2018). We hope our work advances the field by providing a computational framework for mapping theoretical metrics of metacognition onto dynamical models of decision uncertainty.

## Results

### Neural circuit model

Our model comprises two interacting subnetworks. The sensorimotor module comprises two mutually-inhibiting neuronal populations selective for two decision alternatives (eg more dots on the right or left), each of which are endowed with self-excitation (Wong and Wang 2006). Importantly, our model builds on neurophysiological evidence suggesting that decision confidence is encoded by higher-order brain regions (Kepecs et al. 2008; Fleming and Dolan 2012). A crucial aspect of the model is that decision uncertainty (i.e. reciprocal of confidence) is continuously monitored by a dedicated neuronal population termed the ‘uncertainty-monitoring’ population. The latter encodes uncertainty using a leaky integrator – by integrating the summed neuronal activities of sensorimotor populations (see Fig. 1C for a sample trial). Importantly, this integration is terminated when a response is made, i.e. in effect corresponding to when neuronal activity in one of the sensorimotor populations reaches a decision threshold (see Fig. 1C and Methods). Finally, the uncertainty-monitoring population continuously feeds back the encoded uncertainty into both sensorimotor populations via a feedback loop (See Fig. 1B, red arrows). This excitatory feedback mechanism is reminiscent of a dynamic gain modulation (see Fig. 1D), previously shown to account well for response time patterns from decision-making experiments with urgency (Niyogi and Wong-Lin 2013; Smith, Ratcliff, and Wolfgang 2004; Ditterich 2006; Churchland, Kiani, and Shadlen 2008; Kiani, Hanks, and Shadlen 2008; Drugowitsch et al. 2012). Here we refer to this feedback loop as the strength of uncertainty modulation (UM).

**Figure 1.**
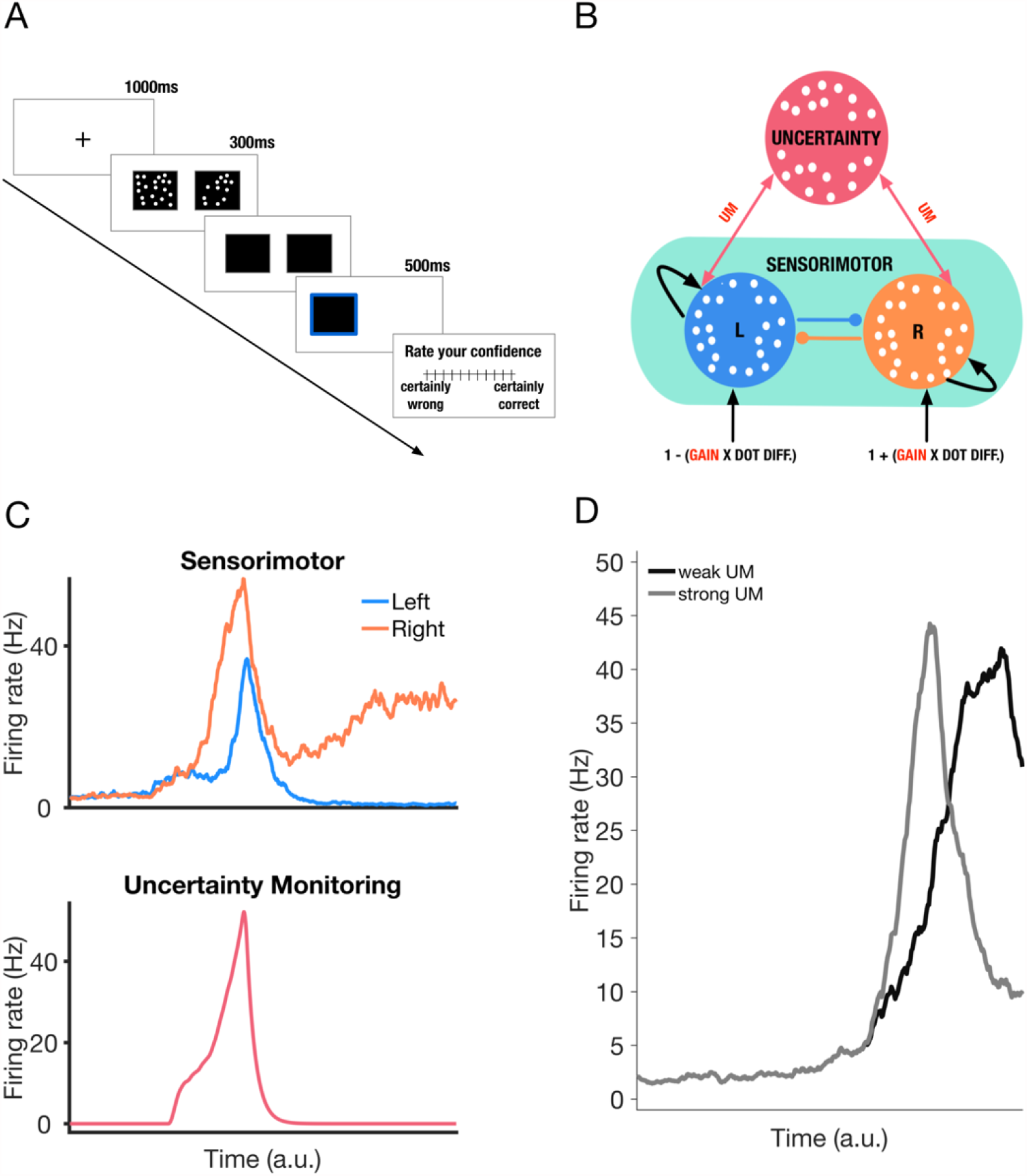
Task and neural circuit model. **A**. Perceptual decision-making task used as a basis for simulations. A fixation cross appears for 1000ms, followed by two boxes with dots for a fixed duration of 300ms. Subjects are asked to judge which box contains the greater number of dots by pressing left/right key on the keyboard. Their response is highlighted for 500ms, i.e. with a blue border appearing around the chosen box. Finally, participants report their confidence in their decision on a scale of 1-11 in experiment 1, and 1-6 in experiment 2 (Supplementary Notes 1 and 2). **B**. Neural circuit model of decision uncertainty. The model comprises two modules. The sensorimotor module (green) comprises two neuronal populations (blue/orange) selective for right/left information. The two populations are endowed with mutual inhibition (lines with filled circles) and self-excitation (curved arrows). These populations receive external input as a function of the difference between the number of dots shown in the two boxes. Figure assumes correct response is on the right – hence the positive input bias for the population selective to rightward information. A gain parameter controls the difference in input each population receives. One neuronal population (red) continuously monitors overall decision uncertainty by integrating the summed output of the sensorimotor populations (see Methods). Uncertainty is equally fed back into both neuronal populations through symmetric feedback excitation (two-way red arrows, controlled by value of uncertainty modulation strength, UM). **C**. A sample timecourse of the activities of the sensorimotor populations (top panel) and uncertainty-monitoring population (bottom panel). Typical winner-take-all behaviour is seen in the sensorimotor module. Activity of the uncertainty-monitoring population follows a phasic profile (see Atiya et al. (2019, 2020) and Methods). Trial simulated with dot difference between the two boxes set at 20 (see Methods). **D**. Sample timecourse of firing rates of the ‘winning’ neural population (i.e. one with more input bias) in the sensorimotor module under two strengths of uncertainty-modulation (UM) values. Random seed reset to control for noise. In the case of the trial with strong (weak) excitatory feedback (solid grey (black) trace), ramping up is faster (slower), leading to a quicker (slower) response. Neural population firing rates shown here are smoothed with a simple moving average (window size = 50ms).

### Applying the model to account for facets of metacognition

We first asked whether our model can account for variation in standard theoretical metrics of metacognition. To do that, we simulated the model using various parameter values, and derived both choices and confidence judgements from the fluctuations in the uncertainty-monitoring population of the model. More specifically, for each simulated trial, we define decision uncertainty (the inverse of decision confidence) as the maximum firing rate reached by the uncertainty-monitoring population within that trial (Atiya et al., 2019). We use equal-width binning to bin (discretise) raw confidence measurements into confidence bins (discrete ratings).

Next, we entered the simulated confidence-accuracy matrix as data into a Bayesian model of metacognitive sensitivity (Fleming 2017). The model returns a parameter *meta – d′* representing the metacognitive sensitivity for a particular simulation with a set of parameter values. Metacognitive efficiency is then estimated by comparing *meta – d′* to the model’s perceptual sensitivity (i.e, *d′*) yielding the ratio meta_d’/d’ (M-Ratio, Maniscalco and Lau 2012). Metacognitive bias is defined as the average binned confidence level across both correct and incorrect trials. We fitted several linear models to estimate the contribution of each parameter in our network model of decision confidence to perceptual sensitivity, metacognitive bias, metacognitive sensitivity, and metacognitive efficiency (see Methods).

The results (Figs. 2A and 2C) show increasing gain has a strong positive effect on *d′* and metacognitive sensitivity. The effect on *d′* is unsurprising given that increasing gain magnifies the difference in input each neuronal population is receiving (see Fig. 1B). *d′* here acts as a ceiling for metacognitive sensitivity, hence the increase in *meta_d’* with increasing gain. Notably, however, we also ran simulations with higher UM values, and metacognitive sensitivity worsened and did not increase with increasing gain, despite the improvement in perceptual sensitivity (Supplementary Note 3). The results also show (Fig. 2B) that increasing gain has a weak effect on metacognitive bias (although see Supplementary Note 4). Finally, the results (Fig. 2D) show that increasing gain has a moderate positive effect on metacognitive efficiency, possibly driven by the sustained linear increase in *d’* as a function of gain.

**Figure 2.**
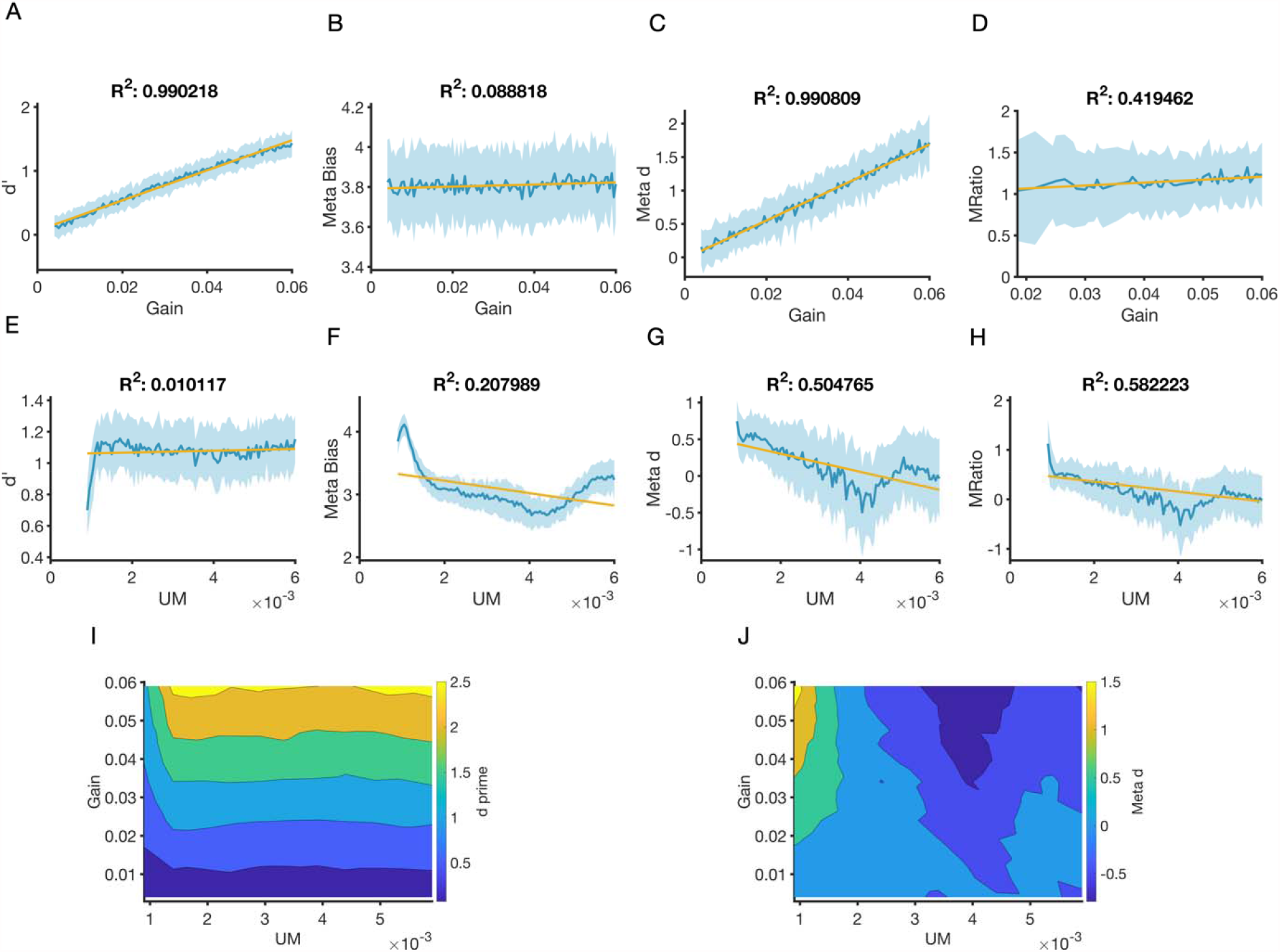
Dissociable changes in metacognition are associated with changes in uncertainty modulation. The behaviour of the model was analysed using standard metrics of performance (d’) and metacognition (metacognitive bias, sensitivity (meta_d’) and efficiency (meta_d’/d’)). Blue line represents mean value of metric across 50 simulations. Shaded area is standard deviation. Yellow line is linear fit to mean value of metric as a function of parameter value. Increases in gain lead to monotonic increases in (**A**) *d′* (*β*_1_ = 0.5, *R*^2^ = 0.99, *p* < 0.001) and (**C**) metacognitive sensitivity (*β*_1_ 28.45, *R*^2^= 0.99, *p* < 0.001) but (**B**) a small effect on bias (*β*_1_ =23.5, *R*^2^ = 0.09, *p* < 0.001). Gain has a moderate positive weak negative effect on (**D**) metacognitive efficiency (*β*_1_ =3.53, *R*^2^ = 0.41, *p* <0.001), possibly driven by the strong linear increase in d’ in panel A. Increasing UM has no effect on (**E**) *d*′(*β*_1_ = 5.65, *R*^2^ = 0.01, *p* <0.15), but a negative effect on (**F**) metacognitive bias (*β*_1_ = -122.83, *R*^2^ = 0.2, *p* < 0.001), (**G**) metacognitive sensitivity (*β*_1_ = -98.83, *R*^2^ =0.5, *p* <0.001), and (**H**) metacognitive efficiency ((*β*_1_ =100.49, *R*^2^ = 0.58, *p* < 0.001). In (**I**-**J**), we varied both parameters and measured the effect on (**I**) *d*′ and (**J**) metacognitive sensitivity. The increase in *d’* is mostly driven by changes in gain (**I**), whereas changes in metacognitive sensitivity are mostly driven by UM (**J**). All simulations were done with the same fixed list of dot differences (2.8 in log-space). In simulations (**A-H**), where the gain (UM) parameter is varied, UM (gain) was fixed at 0.0009 (0.0029). *R*^2^ in all panels is adjusted *R2*. Confidence data was generated by binning the uncertainty values into 6 bins, assuming equal bin width (see Methods). See Supplementary Figure 12 for additional simulations with different parameter values. See also Supplementary Note 5 for results that highlight dissociable changes in metacognitive bias as a result of varying UM.

More interestingly, the second set (Fig. 2, bottom row) of results show that increasing UM has only weak effects on first-order task performance (*d′*) (Fig. 2E). However, increasing UM strength has a negative effect on both metacognitive bias (Fig. 2F) and *meta – d′* (Fig. 2G), leading to reductions in overall confidence and metacognitive sensitivity. Given that first-order performance is relatively unchanged, greater UM strength also results in lower metacognitive efficiency (Fig. 2H). We then varied both parameters together and confirmed that changes in first-order task performance (*d′*) (Fig. 2I) are driven by changes in gain, whereas changes in metacognitive sensitivity (*meta – d′*) (Fig. 2J) are driven by changes in UM.

The bottom row of Fig. 2 suggests a linear fit is not sufficient to account for the relationship between UM and *d’*, meta bias, *meta_d’*, and *meta_d’/d’*. More specifically, despite *d’* remaining mostly constant (∼1-1.1) in the majority of the explored UM parameter space (Fig. 2E), Fig. 2F shows that when UM is between 0.001 and 0.0015, *d’* increases when increasing UM. Furthermore, Figs. 2F, 2G, and 2H show some inflection points where the behaviour before and after these points is different. For instance – meta bias peaks when UM=0.0015, whereas *meta_d’* and *meta_d’/d’* peak at UM=0.005. The existence of nonlinear relationships between UM and our theoretical measures of metacognition is not surprising given that the value of UM governs how two highly non-linear subnetworks of our model interact to generate decision performance and confidence (via increasing/decreasing excitatory feedback).

We note that the model’s uncertainty does not in itself discriminate between correct and incorrect responses. In the model, such differences in confidence naturally emerge through the differences in response times for correct/incorrect trials as a function of difficulty. More specifically, giving the uncertainty-monitoring population more time to integrate input naturally leads to higher uncertainty (less confidence), which in turn is more likely to occur both during incorrect trials and on more difficult problems.

Overall, the results suggest that, in our model, a dissociable uncertainty-monitoring mechanism can drive changes in metacognition, in the absence of any change in task performance. More specifically, stronger uncertainty modulation is associated with a decrease in metacognitive sensitivity, bias, and efficiency, but not perceptual sensitivity. Armed with this understanding of how model parameters relate to facets of metacognitive performance, we next fit the model to subjects’ data, and apply a computational psychiatry approach in order to relate variation in model parameters to psychopathology.

### Model fits to subject data

We re-analysed data from Rouault et al. (2018), in which subjects (experiment 1: 498 subjects, experiment 2: 497 subjects) completed an online task via Amazon Mechanical Turk. In the task, upon initiating a trial, a fixation cross appears for 1000ms, followed by two black boxes each filled with a number of white dots (see Fig. 1A). Subjects indicated first which box contains the greater number of dots, by pressing the right or left arrow key on a computer keyboard, and then provided their confidence rating on a numerical scale (1-11 for experiment 1, 1-6 for experiment 2).

To provide insight into the interaction between decision formation and metacognitive processes in this task, we simulated and fitted our neural circuit model of decision uncertainty to subjects’ choices and response times (Atiya et al. 2019, 2020). This allowed us to use subjects’ explicit confidence reports as an out-of-sample test of the model’s ability to account for individual differences in metacognition. For simplicity, we only simulated the sensorimotor and uncertainty modules of the circuit, as originally introduced in Atiya et al. (2019, 2020) (See Fig. 1B).

In fitting our model to subjects’ choices and response times, we used a procedure based on the subplex optimisation method (Bogacz and Cohen 2004; Rowan 1990) (see Methods). The subplex optimisation method is an evolution of the simplex method (Nelder and Mead 1965) – one that is better suited for optimising noisy objective functions. Importantly, when parameterising our model, we initially set the values of all parameters to those found in our previous work (Atiya et al. 2020), allowing only two parameters to vary in the fitting procedure. The first parameter is a ‘gain’ parameter, which maps the dot difference to input current flowing into the sensorimotor populations (see Methods). Subjects having larger values for the gain parameter generally have better choice accuracy, i.e., at the circuit level, a larger gain value implies a larger bias in sensory input to the sensorimotor population corresponding to the correct choice. The second parameter is the strength of uncertainty modulation (see Fig. 1D for an example of effect of varying this parameter on the decision process).

In experiment 1, subjects completed a perceptual decision-making task in which they judged which box contained a greater number of dots, followed by a confidence report on an 11-point numerical scale. Subjects then completed a number of questionnaires to assess self-reported psychiatric symptoms (see Methods). Unsurprisingly, subjects were more accurate when the task was easy, i.e. when the difference between the number of dots was large (see Fig. 3A). The model captures this straightforward relationship between accuracy and task difficulty (Fig. 3A), and accounts for individual variation in accuracy levels (Fig. 3C).

**Figure 3.**
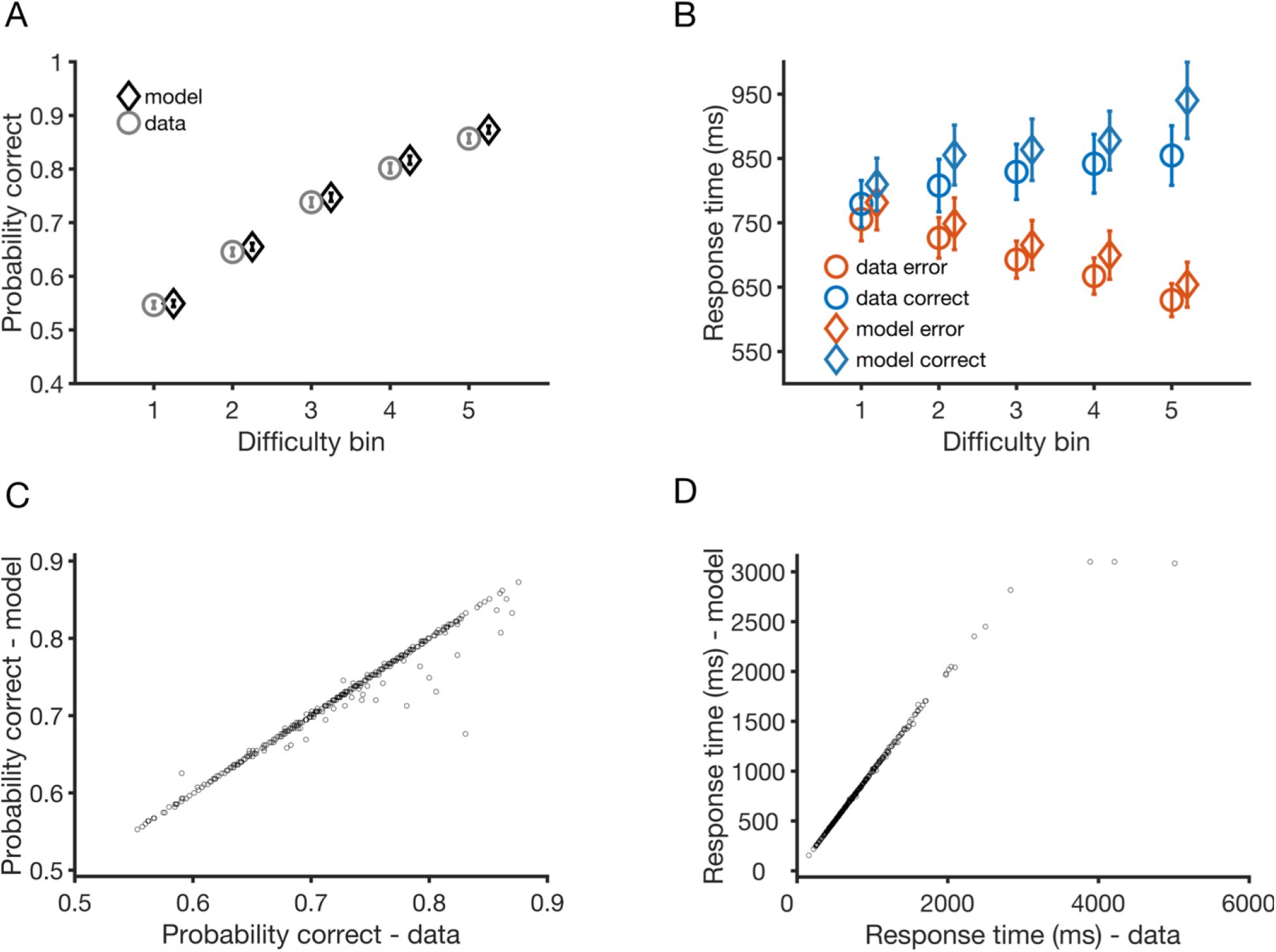
Model accounts for subjects’ perceptual performance in experiment 1. **A**. Choice accuracy, i.e. probability correct as a function of task difficulty from experiment 1 of Rouault et al. (2018) averaged across all 498 participants. Task difficulty is split into 5 difficulty bins (1: most difficult, 5: easiest) as in the original paper (see Methods). Grey markers: data. Black markers: model fits. **B**. Response times as a function of task difficulty from the data (circles) and model fits (diamonds) averaged across all participants. Orange (blue) markers: Error (correct) responses. The typical ‘< ’ pattern, i.e. response times for correct (error) responses increasing (decreasing) as a function of task difficulty, is found in both the model and data. **C**. Scatter plot of observed (empirical) vs. simulated overall accuracy and **D**. response times for each of the 498 subjects. Error bars indicate 95% confidence interval. Random seed is reset after each simulation during the fitting procedure and for the purposes of generating Figures **C** and **D** (but not **A** and **B**). See Supplementary Figure 11 for scatter plots without resetting the random generator seed.

In line with existing findings from both human and animal studies of decision-making (Shadlen and Newsome 2001; Roitman and Shadlen 2002; Sanders, Hangya, and Kepecs 2016), subjects’ correct (error) responses were quicker (slower) as the task became easier, forming a ‘<’ pattern of response times as a function of difficulty (see Fig. 3B, and Fig. 3D for individual variation in mean response time). Observing an interaction between difficulty and accuracy in response time data is particularly striking given that the task was administered using a web-based platform, where response time measurement might be expected to be noisier than in standard laboratory settings. However, such a pattern was closely mirrored by our model fits, and importantly allowed us to constrain the model’s estimates of subjects’ confidence (see below).

### Neuronal model constrained with perceptual performance accounts for subjects’ confidence reports

We next asked whether our fitted model parameters could account for subjects’ explicit confidence reports, even though these data had not been used to constrain the model. Here, we leverage the close relationship between confidence, response time and task difficulty to make inferences about trial-by-trial uncertainty (or confidence) levels from model fits to first-order performance (Kepecs et al. 2008; Kiani, Corthell, and Shadlen 2014). In our model, longer response times allow more time for the uncertainty monitoring population to activate — leading to higher uncertainty (see Methods).

We first simulated our neural circuit model with the parameters fitted to subjects’ choices and response times from experiment 1. We then applied distribution matching (Sanders, Hangya, and Kepecs 2016) to map the model’s simulated uncertainty levels onto subjects’ retrospective confidence reports. More specifically, instead of equal-width binning used in our analyses thus far, the shape of the overall mapping (i.e. prior to conditioning on performance or difficulty) is inferred from the distribution of experimental confidence reports, per subject (see Methods). This allowed us to show the model accounts for the complex relationship between decision confidence and task difficulty (see Fig. 4A). The results also hold after conditioning confidence reports on trial outcome (i.e. correct vs. error). Importantly, these effects result from the intrinsic nonlinear dynamics of the network after fitting to (and constraining the model with) subjects’ first-order performance data alone. The empirical confidence data are only used to set confidence thresholds, prior to conditioning on stimulus difficulty and accuracy. Hence the model is able to account for individual differences in subjects’ perceptual and metacognitive performance despite model fits only having access to choices, response times and the overall distribution of confidence ratings. We next asked whether the uncertainty-monitoring mechanism in the model might also covary with psychiatric symptom scores.

**Figure 4.**
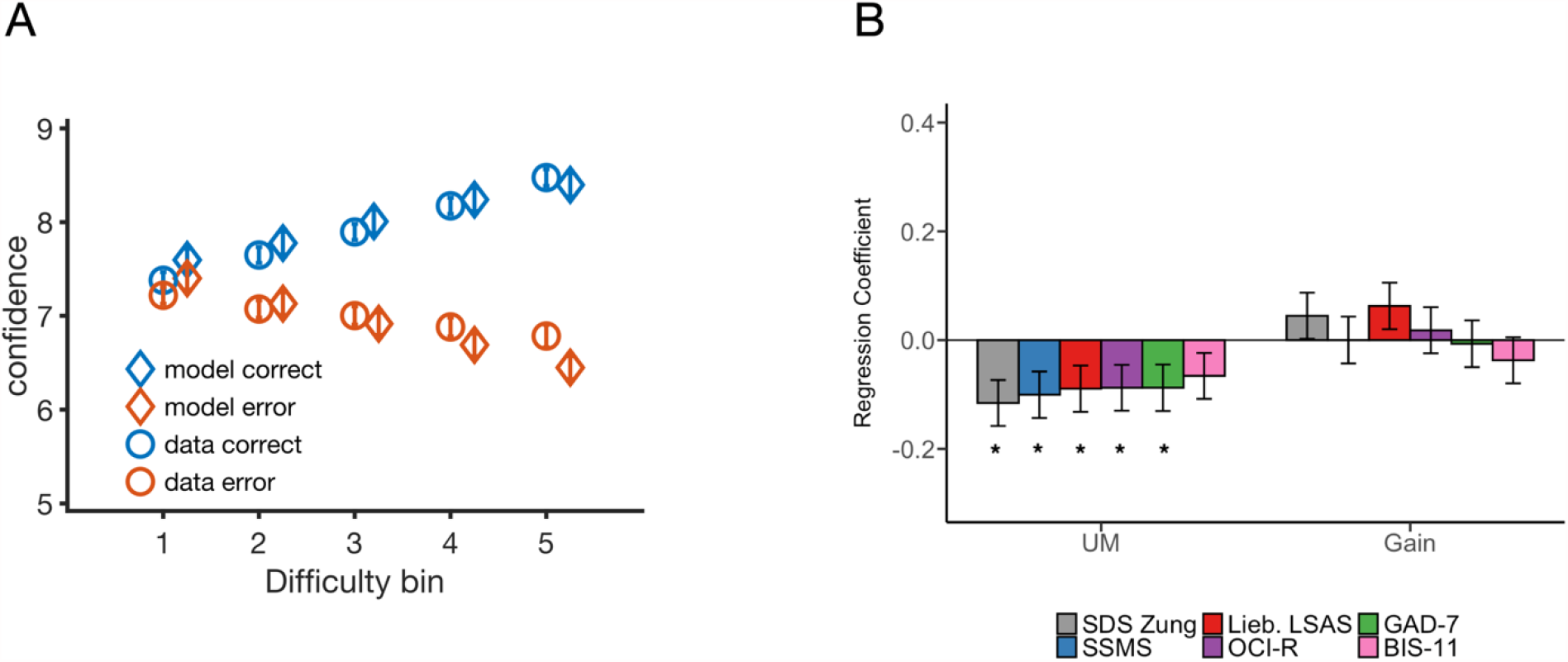
Model accounts for subjects’ confidence reports and individual differences in uncertainty modulation predict symptom scores. **A**. Confidence reports averaged across all participants from experiment 1 data (circles) and model (diamond) as a function of task difficulty. Orange (blue) markers: Error (correct) responses. Note that the model was fit only to first-order performance data (accuracy and response times) and fits to confidence represent an out-of-sample prediction. Confidence increases (decreases) as a function of changing task difficulty for correct (error) responses. **B**. Symptom scores from experiment 1 were entered into a multiple regression model predicting the strength of uncertainty modulation and gain parameters from the model fits to task performance (choices and response times). Self-report measures of depression (grey), schizotopy (blue), social anxiety (red), obsessive and compulsive symptoms (purple) and generalised anxiety (green) are significantly associated with weaker uncertainty modulation. No significant association was found between impulsivity (pink) and the strength of uncertainty modulation. No significant association was found between the symptom scores and the gain parameter. See Methods for details on the regression models. Error bars indicate s.e.m. All regression results shown control for the influence of age, gender, and IQ (see Supplementary Figure 9 for regression model results with age and IQ predicting model parameters). * p < 0.05.

### Psychiatric symptoms are associated with the strength of uncertainty-monitoring

In experiment 1, upon completion of the main perceptual task, participants completed a series of standard self-report questionnaires that assess a range of psychiatric symptoms (Zung 1965; Spitzer et al. 2006; Mason, Linney, and Claridge 2005; Patton, Stanford, and Barratt 1995; Foa et al. 2002; Liebowitz et al. 1985; Spielberger and Gorsuch 1983; Saunders et al. 1993; Marin, Biedrzycki, and Firinciogullari 1991; Garner et al. 1982). The questionnaires comprised: Zung Self-Rating Depression Scale, Generalized Anxiety Disorder 7-item scale, Short Scales for Measuring Schizotypy, Barratt Impulsiveness Scale 11, Obsessive-Compulsive Inventory-Revised [OCI-R], and Liebowitz Social Anxiety Scale.

As in Rouault et al. (2018), we ran a series of linear regressions to tease apart the relationship between psychiatric symptoms and model parameters. Importantly, here, we were able to account for differences in perceptual and metacognitive performance using only two model parameters, as highlighted in our model fits above. The first parameter (UM) controls the strength of uncertainty modulation. The second (gain) parameter maps the dot difference subjects see on the screen to difference in input current flowing into the model’s sensorimotor neuronal populations.

We entered each questionnaire score (see Methods) into multiple linear regressions predicting the uncertainty modulation and gain parameters. The results (see Fig. 4B) show that increases in z-scored self-reported scores were broadly associated with weaker uncertainty modulation across all dimensions of psychopathology, with the exception of impulsivity, though the association strengths did not differ between questionnaires. This contrasts with the gain parameter, which did not correlate with any of the self-reported scores (p> 0.05) in experiment 1. These results largely recapitulate the relationships between empirical confidence level and psychiatric symptoms scores (albeit with minor differences in effect sizes) observed in Rouault et al. (2018), but now provide a potential circuit-level explanation for such differences (i.e., a change in the strength of uncertainty modulation).

We also followed the same approach for experiment 2 (see Supplementary Figure Note 2), although here we found no significant association between the majority of self-reported scores (or cross-cutting factors derived from these scores, see Supplementary Figures 3A and 3B) and model parameters. This lack of significance in experiment 2 may reflect the smaller variance in difficulty (due to the staircase procedure) leading to inferences on uncertainty modulation being less constrained by the data (see Supplementary Figure 10). To explore this further, we attempted to recover the parameters fitted to both experiment 1 and 2 data and found that the fits to experiment 1 data were indeed more stable (See Supplementary Figures 7 and 8) – potentially due to the larger variation in task difficulty. We note however that qualitatively, similar symptom scores (e.g. depression, anxiety) that were negatively related to uncertainty modulation in experiment 1 were also negatively related to the uncertainty modulation in experiment 2. In addition, when using the HMeta-d toolbox (Fleming, 2017) to perform a hierarchical regression (Harrison et al., 2020), we obtained a positive association between the strength of uncertainty modulation and metacognitive efficiency in experiment 2 (mean value of µ_β_ = 0.0516, 95% highest density interval = (0.0813, 0.0016), see Supplementary Figure 6).

## Discussion

While self-reported psychiatric symptoms have been shown to be associated with dissociable differences in metacognition, the mechanisms underlying such changes have remained elusive. In this work, using a computational circuit model of decision-making, we show that shifts in metacognition are associated with disturbances in the interaction between decision-making and uncertainty-monitoring networks. Specifically, stronger uncertainty modulation is associated with decreased metacognitive bias, sensitivity, and efficiency. Importantly, changes in uncertainty modulation strength have no effect on perceptual sensitivity. Notably, our model-fitting approach enabled inferences about uncertainty modulation (and, in turn, these facets of metacognition) from fits to first-order performance data alone. The empirical confidence distributions were used to set confidence thresholds alone (via distribution matching), thus influencing the overall mean and shape of the modelled confidence distribution, but not the relation between confidence and features of performance, response time or task difficulty. Nevertheless, the model is able to account for individual differences in subjects’ perceptual and metacognitive performance despite model parameters being adjusted solely on the basis of patterns of choices and response times. When we apply this approach to data from an online perceptual decision task, we find that self-reported psychiatric symptoms are associated with disturbances in uncertainty modulation.

Through a dedicated uncertainty-monitoring population, our model of decision uncertainty captures key features of the neurobiology of metacognition, while remaining sufficiently simple to fit to data. Recent work has shown that long response times are associated with lower confidence for an impending decision (Kepecs et al. 2008; Kiani, Corthell, and Shadlen 2014; Atiya et al. 2020). Our computational model naturally accounts for this phenomenon. More specifically, winner-take all behaviour is less prevalent when the external stimulus input to the network is (or close to) symmetric, i.e. when stimulus information is ambiguous. This high level of competition between the sensorimotor populations prolongs the time taken to reach a decision threshold, and by allowing more time for an uncertainty-monitoring module to integrate bottom-up input results in higher uncertainty. Building on this proposed mechanism, and existing behavioural evidence, our approach allows us to infer metacognitive performance from first-order (i.e. response time) data.

Crucially, we go beyond simply relating our model dynamics to decision confidence (Atiya et al. 2019). By analysing our model’s uncertainty estimates using standard metrics of metacognition, we reveal that stronger uncertainty modulation in such a network has a negative effect on metacognitive bias, sensitivity, and efficiency, while leaving perceptual sensitivity unaffected. More specifically, controlling for task difficulty, our findings reveal that stronger uncertainty modulation (i.e. leading to overall faster responses) leads to deficits in the accuracy of confidence reports – generally leading to lower confidence in correct trials, and higher confidence in errors. It is of interest to note here that such dissociable changes in metacognitive ability, as a result of a (higher-order) disturbance in the strength of uncertainty modulation, finds support in recent neuropsychological work. For instance, lesions in prefrontal brain regions are associated with deficits in metacognitive ability, but not task performance (Fleming et al., 2014; Lak et al., 2014; Miyamoto et al., 2018), highlighting the contribution of higher-order brain regions to metacognition (Fleming et al. 2010; Fleming and Dolan 2012). Future work could combine our computational framework with neuroimaging to further elucidate the neural basis of metacognitive ability.

Our model architecture complements work exploring how parallel sensorimotor neural populations, with different normalisation tuning, may account for cases where confidence is altered in the absence of a performance change (Maniscalco et al., 2019). We anticipate that a full circuit model of metacognition will need to combine aspects of parallel evidence accumulation (simulating e.g. parietal cortex neural populations) and higher-order monitoring (simulating e.g. prefrontal neural populations involved in the representation and use of uncertainty; Bang & Fleming, 2018). Of particular note here is that we show changes in metacognitive bias may also occur due to shifts in parameters governing higher-order nodes of the circuit. It seems plausible that different forms of confidence bias could map onto different levels of the system. For instance, confidence shifts induced by changes in volatility or amount of evidence may be explained by preferential activation of less-normalisation-tuned populations (Maniscalco et al., 2016), whereas confidence biases related to mood or more “global” aspects of performance may be explained by changes in higher-order nodes of the system (Rouault et al., 2020).

Adopting a computational psychiatry approach, we shed light on a potential driver of metacognitive distortions reported in recent work in relation to mental health symptoms (Rouault et al. 2018). Rouault and colleagues showed that symptom scores for depression, social anxiety, and generalised anxiety relate to lower confidence level. In the present report, following similar analyses, we show that these relationships can be explained by changes in the strength of uncertainty modulation, in the absence of any change in sensory gain. Our analyses not only recapitulate previously-reported relationships with depression and anxiety (Fig. 5B), but show that schizotopy and OCD scores also relate to disturbances in uncertainty modulation (Vaghi et al., 2017), in line with existing work relating deficits in self-evaluation to schizophrenia (Koren et al. 2004).

**Figure 5.**
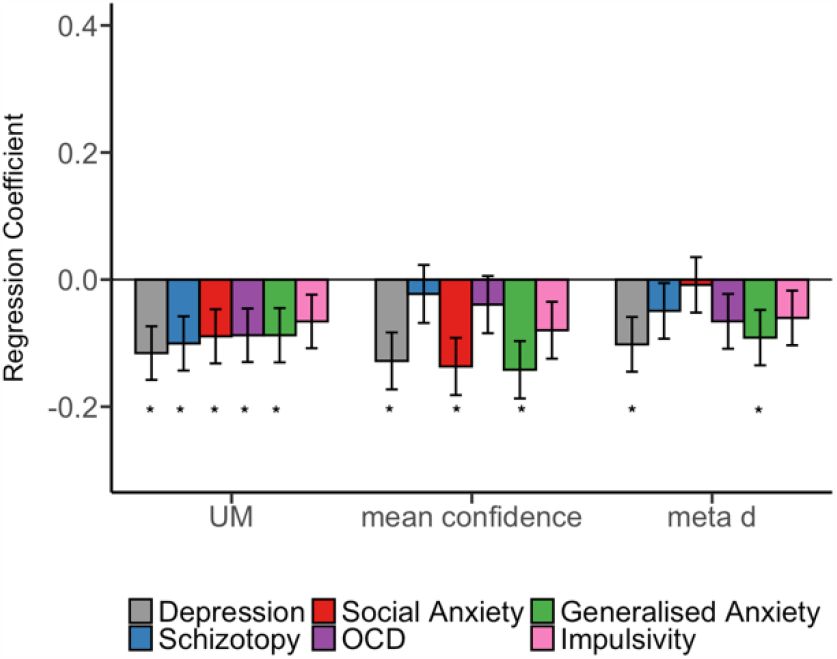
UM parameter offers an implicit, low-dimensional marker of metacognitive (dys)function. Symptom scores from experiment 1 were entered into a multiple regression model predicting the strength of uncertainty modulation, empirical mean confidence, and metacognitive sensitivity. Self-report measures of depression (grey), schizotopy (blue), social anxiety (red), obsessive and compulsive symptoms (purple) and generalised anxiety (green) are significantly associated with weaker uncertainty modulation. Depression, social anxiety, and generalised anxiety are associated with lower mean confidence. Depression and generalised anxiety are associated with decreased metacognitive sensitivity. * p.

In Fig. 5, we compare symptom scores predicting UM (on the left in Fig. 5 below), symptom scores predicting empirical mean confidence (Fig. 5, middle), and symptom scores predicting metacognitive sensitivity (Fig. 5, right). Fig. 5 shows that the standardised effect sizes are slightly larger in the case of Depression (*β* = -0.128, p<0.05), Social Anxiety (*β* = -0.136, p<0.05), and Generalised Anxiety (*β* = -0.141, p<0.05) predicting mean confidence, compared to predicting UM (*β* = -0.115, p<0.05 in the case of Depression, *β* =-0.089, p<0.05 in the case of Social Anxiety, and *β* = -0.087, p<0.05 in the case of Generalised Anxiety). The difference is smaller when comparing the standardised effect sizes in the case of symptom scores predicting UM vs. metacognitive sensitivity (*β* = -0.102, p<0.05 in the case of Depression, and *β* = -0.091, p<0.05 in the case of Generalised Anxiety). Overall, the results suggest that the UM parameter offers an implicit, low-dimensional marker of metacognitive (dys)function, but that confidence rating data would still give a richer perspective and provide distinct measures of sensitivity and bias (as shown in our Supplementary Information Figure 3).

Recent work has demonstrated that symptoms of OCD are associated with deficits in utilising evidence to update confidence (Seow & Gillan, 2020). In the context of our model, this can be explained by the weaker UM strength associated with Obsessive-Compulsive Inventory–Revised (OCIR) scores — i.e. participants with higher OCIR scores tend to monitor uncertainty for longer, prolonging their response times, but not necessarily increasing their confidence in their decisions. Such a mechanism is supported by recent work linking extended evidence accumulation associated with compulsive behaviour to increased decision-making thresholds and metacognitive impairments (Hauser, Moutoussis, et al. 2017; Hauser, Allen, et al. 2017). Notably, in the current work, we could account for individual differences in task (Figs. 3 and 4) and metacognitive performance (Fig. 4A) even in large samples of data (N=495 in Experiment 1, N=496 in Experiment 2 – see Supplementary Note 1 and 2 for Experiment 2 results) collected over the web where experimental control over subjects’ responses is less precise, and response time measurement potentially noisier. Taken together, the results from both experiments suggest our computational framework can be used to study the interaction between metacognition and psychiatric symptoms without requiring subjects to explicitly report confidence in decisions — potentially opening the door to using shorter, more engaging tasks such as smartphone games (Brown et al. 2014).

We also explored whether our model accounts for metacognition-psychopathology relationships in a task with staircased difficulty levels (experiment 2 in Rouault et al.). Although our analyses of the UM parameter show a similar pattern to those obtained for metacognitive bias in the original study (Supplementary Figures 3A and 3B), these relationships between factor scores and model parameters did not reach significance. One interpretation of this equivocal result is that effective inference on individual differences in uncertainty modulation strength may require perceptual tasks with systematic variation in difficulty, to enable full coverage of the RT-accuracy-difficulty surface (i.e. the < patterns). Importantly, we found that the fit for experiment 2 is not as stable as the fit for experiment 1 (Supplementary Figures 4 and 5). Further theoretical work is needed to determine the effect of per-subject difficulty variance on the ability to infer such model parameters.

There are notable limitations to the scope of the model that deserve further investigation in future work. First, it is worth noting that our previous work (Atiya et al., 2019) showed that our model may produce uncertainty and response time patterns that do not strictly follow the ‘<‘ pattern, e.g. with less pronounced increase (decrease) in response times (confidence) for incorrect trials. Such patterns have been previously reported in empirical data (Kiani et al. 2014; Stolyarova et al., 2019; Adler and Ma 2018; Rausch et al. 2018) when considering a wider range of task types (e.g. free-response tasks), and stimulus durations. Second, in our model, there exists a high positive correlation between the maximum firing rate achieved during the trial for both the losing sensorimotor population and the model’s uncertainty monitoring population. This mechanism may limit the model’s ability to account for settings where high (low) confidence is associated with slow (fast) response times. Future modelling work can investigate fitting the model to data from such settings. Finally, previous work (Pleskac & Busemeyer, 2010) has shown that slower response times can be associated with higher confidence when subjects optimise for accuracy over speed. In our model, both the non-selective top-down excitation of the sensorimotor populations and the decision time (which determines the integration window of the uncertainty-monitoring population) contribute to a high positive correlation between uncertainty and response time. Accounting for a reversal in this relationship is beyond the scope of the current modelling work. In order to account for various speed/accuracy trade-offs leading to differential confidence-reponse time correlations, future work will need to investigate alternative model architectures where the integration-time window is fixed across trials, and is not decision-time dependent.

Previous versions of our neural circuit model have also been applied to tasks with explicit motor reaching trajectories through a dedicated motor output network (Atiya et al. 2019, 2020). Here, given that participants reported their decisions using a keyboard button press rather than continuous motor responses, this aspect of the network was less relevant. However, our current findings highlight the promise of leveraging the full model to dissect the interaction between uncertainty-monitoring, indecisiveness and psychiatric symptoms in a task where both sensory input and motor output are quantified in a continuous, dynamic fashion. Because these relationships can be obtained from fits to first-order performance and response time data alone, future work could leverage our computational framework to infer facets of metacognition in situations where obtaining explicit metacognitive judgements is problematic or impossible, e.g. in studies of animals or children.

In summary, we employed a biologically-plausible model of decision uncertainty to relate dissociable shifts in metacognition to isolated disturbances in uncertainty modulation. We validate our model against empirical data, and relate its parameters to psychopathology.

Our work bridges a gap between a biologically plausible model of confidence formation and the observed disturbances in metacognition seen in mental health disorders, and provides a first step towards mapping theoretical constructs of metacognition onto dynamical models of decision uncertainty. In doing so, we provide a computational framework for modelling metacognitive performance in settings where access to explicit confidence reports is either difficult or impossible.

## Methods

### Neural circuit model of uncertainty

We modelled the processes underpinning decisions and confidence using a neural circuit model of uncertainty described previously (Atiya et al. 2019, 2020). The version of the model used here comprises two interacting subnetworks — a decision-making *sensorimotor module*, and an *uncertainty-monitoring* population.

As in previous work (Atiya et al. 2019, 2020), the sensorimotor module is modelled using a reduced (i.e. two-variable) spiking neural network model (Wang 2002; Wong and Wang 2006). The dynamics of the neuronal populations are described by:

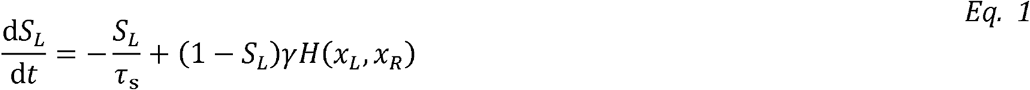

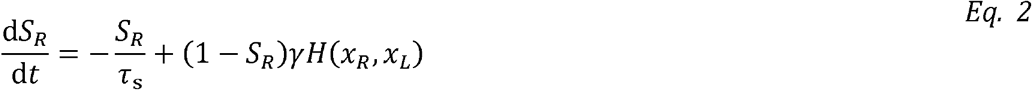

where *S*_*L*_ and *S*_*R*_ are the synaptic gating variables for the sensorimotor population selective to leftward and rightward stimulus information, respectively. *τ*_s_ denotes the synaptic gating time constant. *γ* is a constant that is derived in previous theoretical work (Wong and Wang 2006) that describes a reduction of the original spiking neuronal network model of decision making (Wang 2002).

The firing rate of a sensorimotor population can be described using the nonlinear function *H*:

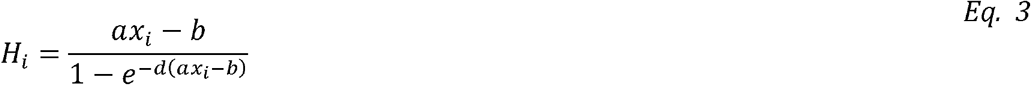

where *a, b, d* are parameters fitted to the leaky integrate-and-fire model (Wang 2002). The variable *i* can be *L* or *R*, denoting sensorimotor population selective for rightward or leftward sensory information, respectively. *x*_*i*_ denotes the total input into population, and can be described by:

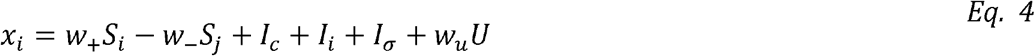

where *w*_+_ denotes synaptic weight for self-excitation, whereas *w*_-_ denotes synaptic weight for mutual inhibition. *I*_*c*_ is some constant input. *I*_σ_ denotes noise — here we use the same noise described by an Ornstein–-Uhlenbeck process as in (Wong and Wang 2006). *I*_*i*_ denotes external input flowing into population *i*, as a function of the dot difference participants see on the screen (Fig. 1). This external input is described by:

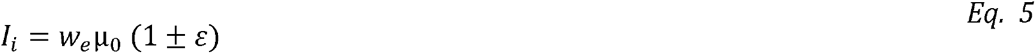

where *w*_e_ is a synaptic weight, whereas *µ*_0_ is some baseline external input. *ε* can be described by:

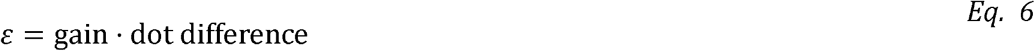

where the input gain parameter maps the dot difference to difference in input flowing into the sensorimotor populations. In our model, sensorimotor populations continue to integrate evidence for 180ms after the initial decision is made (nondecision time). This nondecision time has been used in previous work to account for signal transduction delays (Resulaj et al., 2009; Albantakis et al., 2011). Our model does not account for more extended post-decisional processing, or the incorporation of new, post-decisional evidence. Such additions to the model may usefully augment the extent to which we are able to accommodate dissociations between confidence and performance.

Importantly, the last term in Eq. 4 (*w*_*u*_*U*) determines the strength of feedback excitation from the uncertainty-monitoring neuronal population. More specifically, *w*_*u*_ is referred to throughout this article as UM, or uncertainty modulation strength. U denotes the dynamical variable of the uncertainty-monitoring population, which is described by:

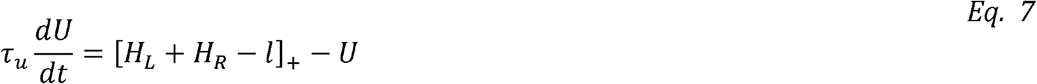

where []_+_ is a threshold linear function (threshold = 0). *H*_*L*_ and *H*_*R*_ are functions denoting firing rates for sensorimotor populations selective for leftward and rightward stimulus information, respectively (from Eqs. 1 and 2). *l* denotes some constant input that suppresses the firing of the uncertainty-monitoring population. This input is de-activated 200ms after stimulus onset, and is reactivated when one the firing rate of the sensorimotor populations reaches a decision threshold (see Fig. 1). Eq. 7 includes a leak term (*–U*), hence why the integration decays over time when no external input is present (i.e., a leaky integrator). We summarise the values of all model parameters in Table 1.

**Table 1.**
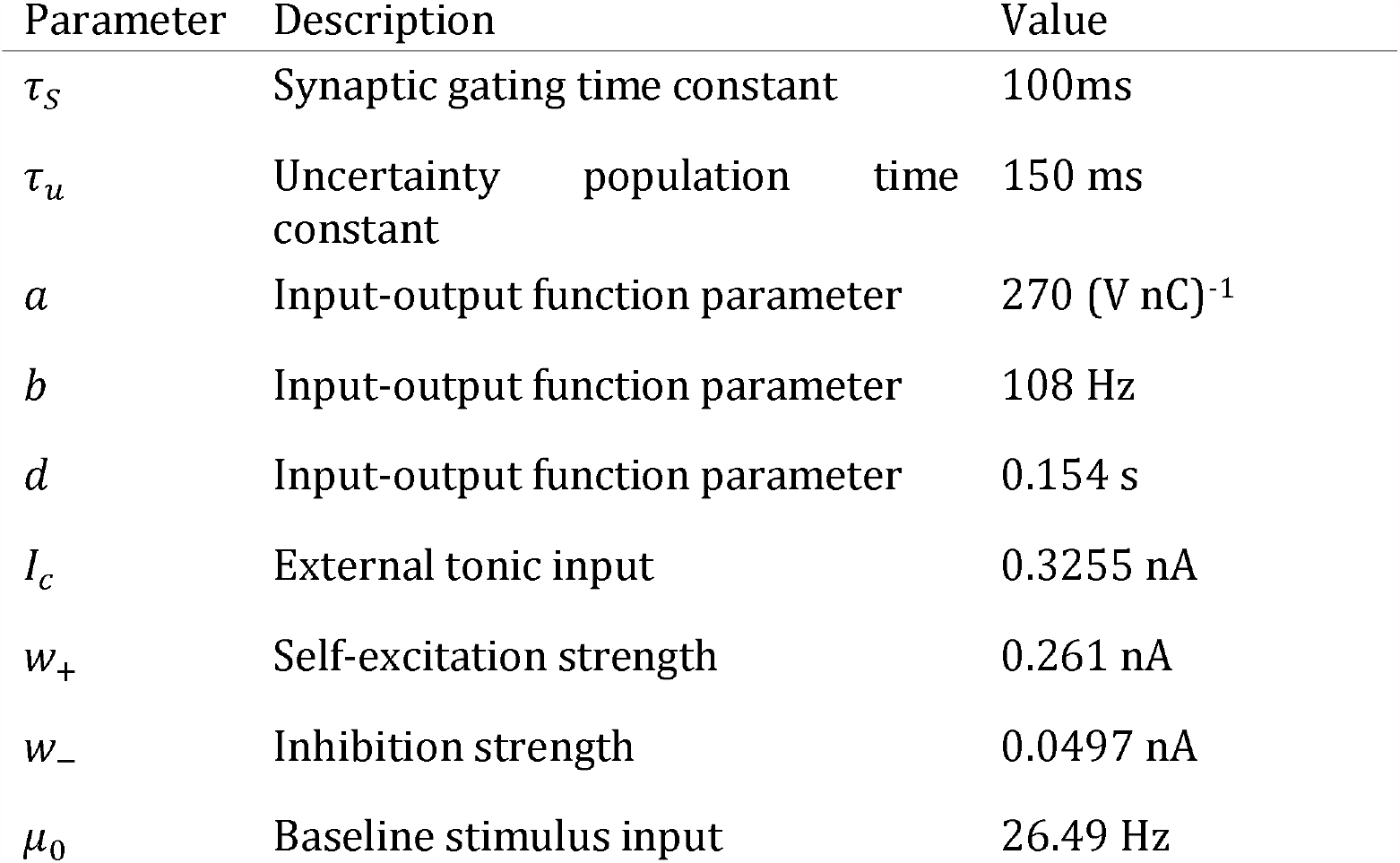
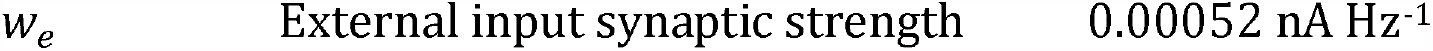
Table of fixed model parameter values for all participants. Parameters *τ*_*S*_, *τ*_*u*_, *a, b, d, I*_*C*_, *w*_*e*_ were directly adapted from (Atiya et al. 2020). Parameters *μ*_*0*_, *w*_*+*_, *S*_*th*_ were manually tuned to adapt the model simulations to the task and stimuli.

### Quantifying uncertainty within a trial

As in our previous work (Atiya et al. 2020), for a given trial, we used the maximum firing rate value of the uncertainty-monitoring neuronal population as a decision uncertainty measurement for that particular trial (the inverse of decision confidence). When extrapolating confidence reports from simulations (e.g. for Fig. 2 simulations), we used simple equal-width binning in 6 bins to relate continuous uncertainty measurements to a 6-point confidence scale, similar to the one used in experiment 2.

Each participant uses the confidence scale differently, e.g. on a 6-point probabilistic scale, one might consistently pick 5 as their highest confidence level. In order to relate simulated uncertainty to empirical confidence data from each participant, we match the distribution of simulated uncertainty to the marginal distribution of empirical confidence reports (i.e. prior to conditioning on accuracy, response times, or difficulty; Sanders, Hangya, and Kepecs 2016). More specifically, per subject, we (non-parametrically) infer the shape of the mapping from their experimental confidence distribution. First, we compute the cumulative distribution function (CDF) of their full confidence distribution. Then, we use this CDF to derive binning width thresholds. The thresholds here represent the quantiles of the subjects’ simulated confidence for the probabilities represented by CDF computed from experimental confidence distribution.

### Model fitting procedure

To fit our model to participants’ first order performance, we used a procedure that exploits the subplex optimisation method (Bogacz and Cohen 2004; Rowan 1990). Subplex optimisation is based on the simplex optimsation method, but adapted for noisy objective functions (Rowan 1990). For each participant, we minimise the cost function:

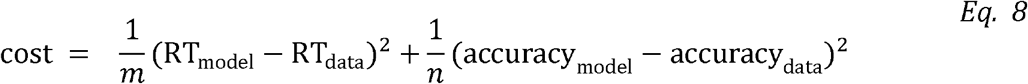

where RT_model_ is the model’s mean response time from a single model simulation (with a fixed random seed), RT_data_denotes the participants’ mean response time. Similarly, accuracy_model_ and accuracy_data_ denote overall accuracy for the model and experiment, respectively. *m* and *n* are normalisation terms for response times and accuracy, respectively. Here, *m* and *n* are set to the model statistic (i.e., *m* = RT_model_, and *n* = accuracy_model_) (Bogacz & Cohen, 2002). The cost function can be calculated per difficulty level (see Berlemont et al. (2020)). Here, we opted for calculating the cost using the overall accuracy and overall response times (across all difficulties). Importantly, we only fit two free parameters: gain and w_u_, from Eqs. 6 and 4, respectively. The vast majority of the other model parameters are adapted from our previous work (Atiya et al. 2020) (see Table 1). When generating synthetic data using the model (for fitting or otherwise), for experiment 1, we simulate 210 trials while generating dot difference data from a uniform distribution bounded by the max and min value for each difficulty block as found in the data. For experiment 2, we simulate the model with the vector of dot differences experienced by each participant.

### Ethics statement

Data analysed in this work was first collected as part of a study conducted by Rouault et al. (2018). Participants provided written consent in accordance with procedures approved by the University College London Research Ethics Committee (Project ID 1260/003).

### Participants

We re-analysed data from Rouault et al. (2018), and the reader is referred to this paper for a full description of the task and sample. All participants were recruited over the web using Amazon Mechanical Turk. In experiment 1, 663 (498 after exclusions) participants completed the task, and were 18-75 years of age. In experiment 2, 637 (497 after exclusions) participants completed the task, and were 18-70 years of age. The study protocol was approved by the University College London Research Ethics Committee (REF 1260/003) and all participants provided informed consent before undertaking the task. All participants in experiment 1 and 2 were compensated $4. A $2 bonus was paid out to participants on two conditions: In experiment 1, the bonus was paid if participants achieved >50% accuracy in task performance, and passed a check question. In experiment 2, the bonus pas paid if participants achieved task performance between 60-85%, and passed a check question. We used the same exclusion criteria applied in Rouault et al. (2018) and described in the Supplementary Material of that paper.

### Task

In both experiments, participants completed a simple perceptual decision-making task where they judged (using a keyboard press) which box contained a higher number of dots, with no feedback. One box was always half-filled (313 dots out of 625 positions), while the other box contained an increment of +1 to +70 dots compared to the standard. In any given trial, a fixation cross first appeared for 1 second, followed by two black boxes with two different amounts of dots (for 300ms). The position of the box with higher number of dots (i.e. target box) was pseudo-randomised. After indicating the position of the target box (left/right) via a keyboard arrow button press, the box was highlighted for 500ms. In experiment 1, participants completed 210 trials, split over 5 blocks, where the difficulty was varied. The position of the target box was pseudo-randomised across all trials and within each of 5 difficulty bins.

After every trial, participants provided a confidence judgement on a full 11-point probabilistic scale (Boldt and Yeung 2015): 1=certainly wrong, 3=probably wrong, 5=maybe wrong, 7=maybe correct, 9=probably correct, 11=certainly correct. Finally, pre- and post-task global confidence ratings were given by participants, together with their estimates of expected maximum and minimum levels of task performance.

Prior to undertaking the experiment, participants were required to select on an 11-point scale their global expected performance level in the task relative to others, together with a maximum and minimum expected performance level. After completing the task, participants were again asked to rate their expected performance level in the task relative to others, using the same scale. Pre- and post-global confidence levels were not analysed here.

Experiment 2 (see Supplementary Note 1) is identical to experiment 1 in all but three aspects. First, Rouault et al. (2018) used a staircase (calibration) procedure to fix participants’ perceptual performance (Garcia-Pérez 1998; Fleming et al. 2010). The staircase procedure was two-down one-up, with equal step sizes. Step-sizes (in logspace) were: 0.4 for first 5 trials, 0.2 for next 5, 0.1 for the rest of the task. The starting point was 4.2. Each participant completed 25 practice trials at the beginning of the task to minimise the burn-in period. Second, participants reported their confidence on a 6-point confidence scale which ranged from 1= guessing to 6=certainly correct). Third, pre- and post-task global confidence ratings were omitted from experiment 2.

The entire experiment was coded in JavaScript with JsPsych version 4.3 (De Leeuw JR, 2015).

### Psychiatric questionnaires

Participants completed a set of self-report questionnaires used to assess their psychiatric symptoms (Rouault et al. 2018). In experiment 1, the questionnaires were:

- Depression using the Self-Rating Depression Scale (SDS) (Zung 1965).
- Generalised anxiety using the Generalised Anxiety Disorder 7-Item Scale (GAD-7) (Spitzer et al. 2006)
- Schizotypy using the Short Scales for Measuring Schizotypy (SSMS) (Mason, Linney, and Claridge 2005)
- Impulsivity using the Barratt Impulsiveness Scale (BIS-11) (Patton, Stanford, and Barratt 1995)
- Obsessive Compulsive Disorder (OCD) using the Obsessive-Compulsive Inventory– Revised (OCI-R) (Foa et al. 2002)
- Social anxiety using the Liebowitz Social Anxiety Scale (LSAS) (Liebowitz et al. 1985)

In experiment 2 (Supplementary Notes 1 and 2), the following changes were made to the set of questionnaires:

- Generalised Anxiety questionnaire was replaced by the State Trait Anxiety Inventory (STAI) Form Y-2 (Spielberger and Gorsuch 1983)
- Alcoholism was assessed with the Alcohol Use Disorders Identification Test (AUDIT) (Saunders et al. 1993)
- Apathy was assessed with the Apathy Evaluation Scale (AES) (Marin, Biedrzycki, and Firinciogullari 1991)
- Eating disorders was assessed with the Eating Attitudes Test (EAT-26) (Garner et al. 1982)

These changes in experiment 2 were made to facilitate identification of three latent factors that accounted for the majority of covariance across individual questionnaire items (Gillan et al. 2016).

### Factor analysis

For experiment 2 data (see Supplementary Notes 1 and 2), we obtained three latent factors that explain the shared variance across the 209 questionnaire items. To do that, we followed the same approach in Rouault et al. (2018) and Gillan et al. (2016), and used the *fa()* function from the Psych package in R. The three latent factors were Anxious-Depression, Compulsive Behaviour and Intrusive Thought, and Social Withdrawal.

### Linear regressions

To estimate the relationship between the neural model parameters and self-reported psychiatric scores, we followed the same approach as in Rouault et al. (2018). All regressors were z-scored to ensure comparability of regression coefficients. For each symptom score, and controlling for age, IQ and gender the regressions were:

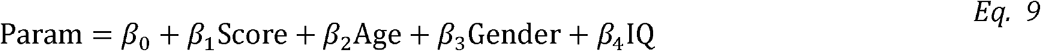

To assess the relationship between model parameters and the latent factor scores (see above), the regression was:

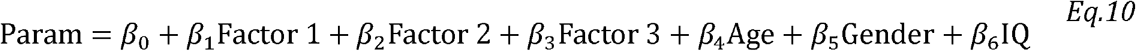

Finally, we used linear regressions to estimate the contribution of two of the model parameters to standard metrics of metacognition and perceptual sensitivity. Here, we did not z-score the regressors as the goal was to visualise the relationship rather than quantitatively compare coefficients. The regressions were:

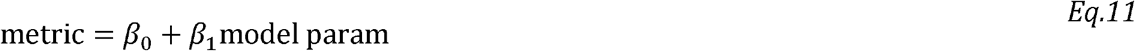

### Metacognitive bias, sensitivity, and efficiency

Metacognitive bias was computed as the mean confidence level across both correct and incorrect trials. To estimate metacognitive sensitivity, we entered simulated confidence reports as data in a Bayesian model of metacognitive efficiency, HMeta-d (Fleming 2017). The model returns a value of metacognitive sensitivity (*meta – d′*) for each simulated dataset. To compute metacognitive efficiency, we calculated the ratio *meta – d′/d′*.

## Data availability

Code used to fit, simulate, and analyse the model (and data) is available at this repo: https://github.com/nidstigator/uncertainty_psychiatry_2020

Data collected is available in the same repo.

## Acknowledgments

SMF is funded by a Wellcome/Royal Society Sir Henry Dale Fellowship (206648/Z/17/Z) and a Philip Leverhulme Prize from the Leverhulme Trust. The Wellcome Centre for Human Neuroimaging is supported by core funding from the Wellcome Trust (206648/Z/17/Z). The Max Planck UCL Centre is a joint initiative supported by UCL and the Max Planck Society. The funders did not play any role in the study design, data collection and analysis, decision to publish, or preparation of the manuscript.

## Supporting Information Captions

**S1 Note. Model fit to experiment 2 data from Rouault et al. (2018)**.

**S1 Figure. Model accounts for subjects’ perceptual performance in experiment 2. S2 Figure. Model accounts for subjects’ confidence reports in experiment 2**.

**S2 Note. Relationships between psychiatric symptoms and model parameters in Experiment 2**.

**S3 Figure. No significant relationships between psychiatric symptoms scores and model parameters**.

**S3 Note. Exclusion Criteria**.

**S4 Note. Integration onset timing parameter**.

**S5 Note. Changes in metacognitive bias driven by UM**.

**S4 Figure. Changes in metacognitive bias are driven by changes in parameter governing higher-order nodes of the circuit (UM parameter) independently of changes in performance (d′)**.

**S5 Figure. Model parameters fitted to a subset (70%) of the data were then used to simulate data for each participant**.

**S6 Figure. Hierarchical estimation of the impact of fitted UM on observed meta_d’/d’ ratio using a simultaneous regression approach with the UM parameter as a covariate (Harrison et al**., **2020)**.

**S6 Note. Experiment 1 fit with holdout**.

**S7 Note. UM as a low dimensional marker of metacognitive profile**.

**S7 Figure. Results from parameter recovery simulations for experiment 1 for both (A) Gain and (B) UM**.

**S8 Figure. Results from parameter recovery simulations for experiment 2 for both (A) Gain and (B) UM**.

**S9 Figure. Relationship between Age/IQ and model parameters**.

**S10 Figure. Variance in difficulty experienced by participants in Experiment 1 and 2**.

**S11 Figure. (A) Individual accuracy and (B) mean response time model fits for Experiment 1 without resetting the random number generator seed**.

**S12 Figure. Simulations of Figure 2 in the main manuscript with alternative parameter values**.

