## Supplementary Information for "Explaining distortions in metacognition with an attractor network model of decision uncertainty"

**Supplementary Note 1: Model fit to experiment 2 data from Rouault et al. (2018)**

Experiment 2 (n=497) is identical to experiment 1 in all but three aspects. First, Rouault et al. (2018) used a staircase (calibration) procedure to fix participants’ perceptual performance (Garcıa-Pérez 1998; Fleming et al. 2010). The staircase procedure was two-down one-up, with equal step sizes. Step-sizes (in logspace) were: 0.4 for first 5 trials, 0.2 for next 5, 0.1 for the rest of the task. The starting point was 4.2. Each participant completed 25 practice trials at the beginning of the task to minimise the burn-in period. Second, participants reported their confidence on a 6-point confidence scale which ranged from 1= guessing to 6=certainly correct). Third, pre- and post-task global confidence ratings were omitted from experiment 2.

Similar to our analysis of Experiment 1, we first fitted our neural circuit model to subjects’ choices and response times, but not confidence reports (see Methods). The results show (see Supplementary Figs. 1A and 1B) that our model again accounts for both average patterns of choice accuracy and response times, and individual differences across participants (Supplementary Fig. 1C and 1D).

To fit the model to subjects’ confidence reports, we first simulated our neural circuit model with the parameters fitted to subjects’ choices and response times from experiment 2, and applied distribution matching to map the model’s simulated uncertainty levels onto subjects’ retrospective confidence reports (see Methods in main text). The results (Supplementary Figure 2) show that the model accounts for the complex relationship between decision confidence and task difficulty, and the results hold after conditioning confidence reports on the outcome of the trial (i.e. correct vs. error).

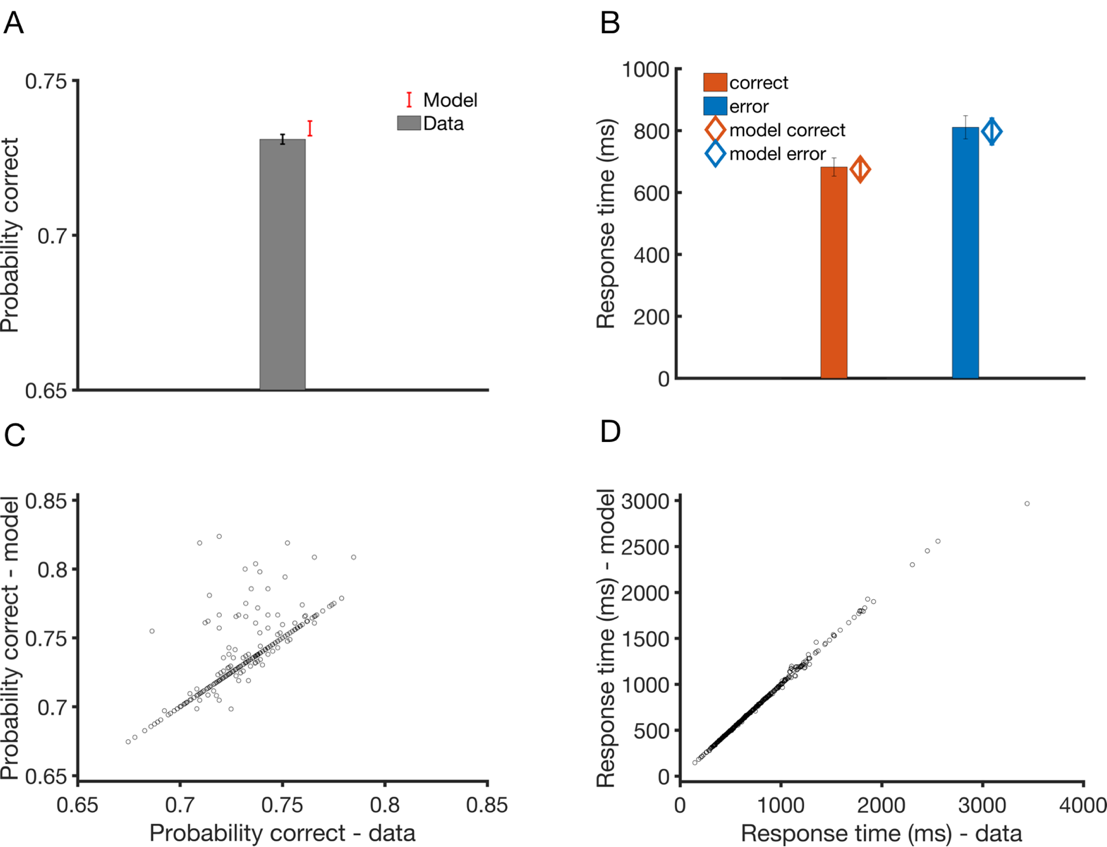

**Supplementary Figure 1.** **Model accounts for subjects’ perceptual performance in experiment 2**. **A.** Choice accuracy averaged across all participants from experiment 2. Model fit (red) accounts for the data (grey bar). **B.** Response times averaged across all participants from experiment 2, split by correct (orange) and error (blue) responses. Slower (faster) responses in case of error (correct) responses are observed both in the model (markers) and data (bars). Error bars indicate 95% confidence interval. **C.** Scatter plot of observed vs. simulated mean response times and **(D)** accuracy for each of the 497 subjects. Random seed is reset after each simulation during fitting procedure and for the purposes of generating figures C and D.

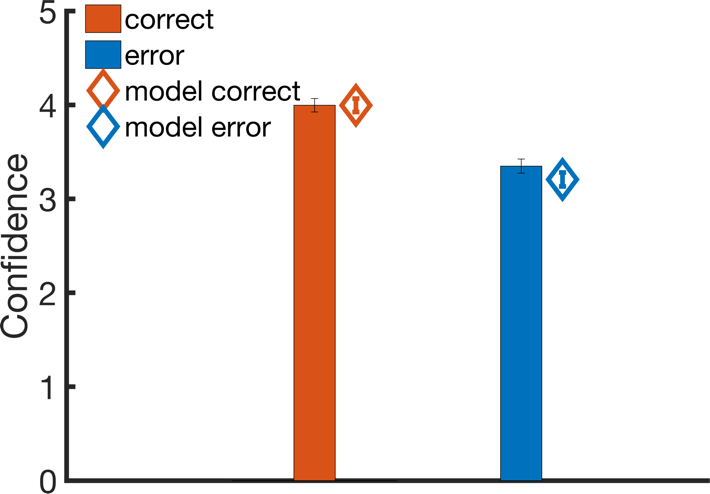

**Supplementary Figure 2. Model accounts for subjects’ confidence reports in experiment 2.** Confidence reports averaged across all participants from experiment 2 data (bars) and model (markers). Blue (orange) markers: Error (correct) responses. Error bars indicate 95% confidence interval.

**Supplementary Note 2: Relationships between psychiatric symptoms and model parameters in Experiment 2**

In experiment 2, additional questionnaires were added (Eating Attitudes Test, Apathy Evaluation Scale, and Alcohol Use Disorders Identification Test), and the Generalized Anxiety Disorder 7-item scale was replaced by the State Trait Anxiety Inventory (STAI) questionnaire. Additionally, participants completed a short IQ test (International Cognitive Ability Resource) (Condon and Revelle 2014).

We found no significant association between self-reported scores (or cross-cutting factors derived from these scores, see below) and model parameters (Supplementary Figure 3). We considered whether this lack of significance in experiment 2 may have been due to the smaller variance in difficulty (due to the staircase procedure) leading to inferences on uncertainty modulation being less constrained by the data than in experiment 1 (see Supplementary Figure 5). To explore this further, we attempted to recover the fitted parameters to both experiment 1 and 2 data and found that the fit to experiment 1 data is indeed more stable (see Supplementary Figures 4 and 5) – potentially due to the difference in difficulty variation (see Supplementary Figures 7). We note however that qualitatively, similar symptom scores (e.g. depression, anxiety) that were negatively related to uncertainty modulation in experiment 1 were also negatively related to uncertainty modulation in experiment 2.

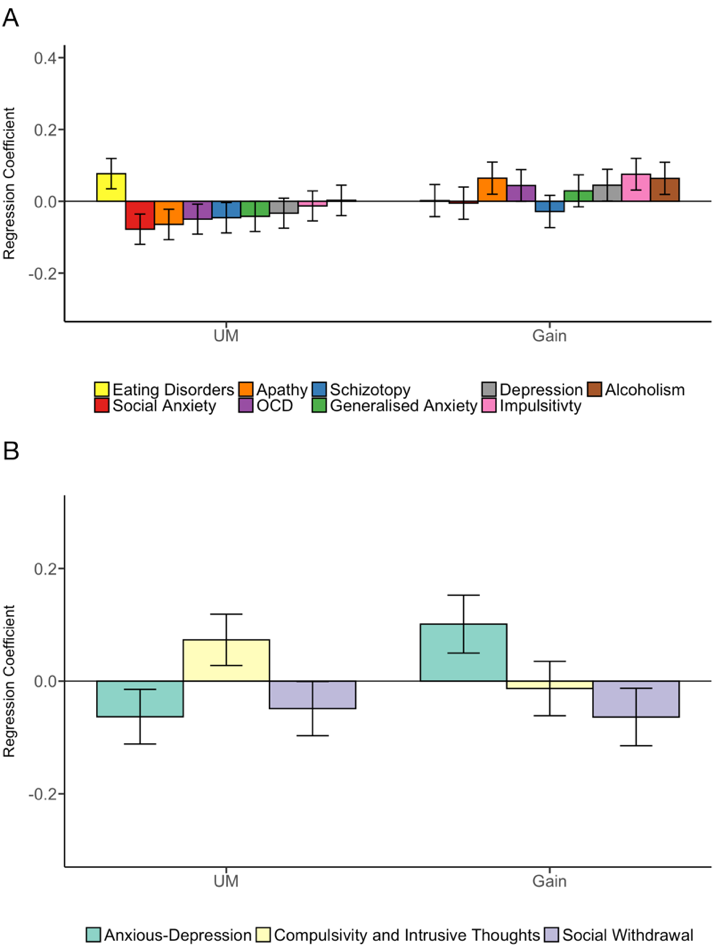

**Supplementary Figure 3. No significant relationships between psychiatric symptoms scores and model parameters.** **A**. Symptom scores from experiment 2 were entered into a multiple regression model predicting the model parameters. No significant association was obtained between individual symptom scores and model parameters, although qualitatively we again observed negative relationships between uncertainty modulation and several of the same symptom scores (e.g. depression and anxiety) that were significantly negative in experiment 1. **B**. Three latent factors based on symptoms scores from experiment 2 (see Methods) were entered into a multiple regression model predicting model parameters. These were Anxious-Depression (green), Compulsivity and Intrusive Thought (yellow), and Social Withdrawal (purple). No significant relationships between the factors and model parameters were observed. See Methods for details on the regression models. Error bars indicate s.e.m. All regressions results shown control for the influence of age, gender, and IQ.

**Supplementary Note 3: Exclusion Criteria**

Experiment 1

Participants were required to pass all of the following 6 tests to be included:

1. Performance of at least 55% correct
2. Prior to the task, a test to measure how well participants understood the confidence scale (what would you rate your confidence if you were sure your judgment was correct (incorrect)? Correct answer is 11 (1)
3. When rating pre- and post- global expected performance level: minimum expected performance level ≤ average performance level ≤ maximum expected performance level
4. A “catch” question: “If you are paying attention to these questions, please select ‘A little’ as your answer”
5. Not repeatedly selecting the same confidence rating across trials

Experiment 2 had the same exclusion criteria adopted – with the addition of prohibiting participants who took part in Experiment 1 from taking part in Experiment 2.

**Supplementary Note 4: Integration onset timing parameter**

In our model, we use an inhibitory mechanism to gate the integration of input in the uncertainty monitoring population. Such an inhibitory mechanism has been proposed to originate from a subcortical circuit. For example, the threshold crossing (response threshold in our model, which triggers top-down inhibition) could be detected by the superior colliculus via basal ganglia (Lo and Wang, Nat. Neurosci. 2006; Crapse and Sommer, J. Neurosci., 2009). More complex gating pathways in the brain, including disinhibitory circuits, have been proposed to also involve subcortical structures, such as the basal ganglia and thalamus (Wang & Yang, Curr. Biol., 2018). As a proxy for modelling such complex and extended neural networks, we instead modelled just the onset and offset of top-down inhibition. The onset of this top-down inhibition is assumed to have been learned e.g. through the basal ganglia (see Hazy et al., Proc. Royal Soc. B, 2007), via changes in the influence of neuromodulators (Frank, J. Cogn. Neurosci., 2005). In this sense, the exact onset time (mediated partially through gating and top-down inhibitory mechanisms) can be learned using models such as the one outlined by Alexander and Brown (Nat. Neurosci., 2011). Providing an explicit account of such complex neural circuit dynamics is beyond the scope of this work, and we hope future work will address this in more detail.

The timing value (200ms) has been explored in detail in our previous work (Atiya et al. 2019). In this work, we decided to fix this value across all subjects and vary only the other two parameters.

**Supplementary Note 5: Changes in metacognitive bias driven by UM**

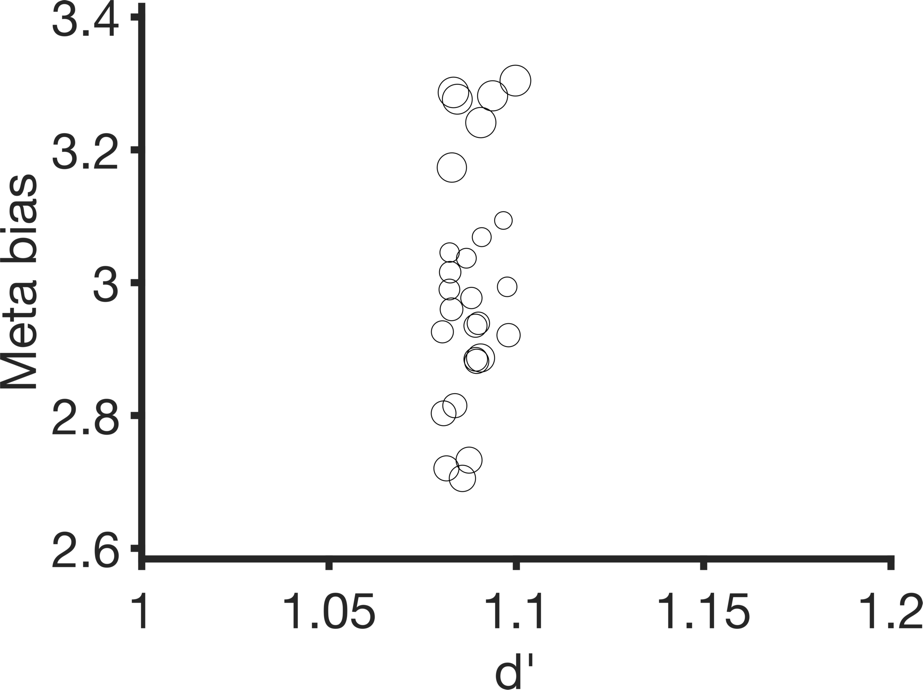

**Supplementary Figure 4.** **Changes in metacognitive bias are driven by changes in parameter governing higher-order nodes of the circuit (UM parameter) independently of changes in performance (**$\boldsymbol{d'}$**)**. Each circle is a mean of 50 simulations. For each circle, the same dot difference (2.8 in log space) and gain parameter (0.025) were used, but UM was varied between 0.002 and 0.006. Size of circle indicates UM value (larger circle = larger value).

More specifically, we simulated our model under a fixed difficulty level (dot difference set at 2.8 in log space), thereby maintaining a constant d’. We then examined the effect of 27 UM different parameter values spaced between 0.02 – 0.06, while fixing the gain parameter at 0.025 (as in our other analyses, e.g. Fig. 2 in the main manuscript). Each simulation was repeated 50 times. Our results show that, in this particular subspace of the model parameters, UM has a strong effect on mean confidence but not d’.

In Maniscalco et al. (2021), the authors reproduce a similar result – matched d’ but introducing changes in confidence (Koizumi et al., 2015) – by computing confidence primarily from the ‘less normalised units’ in their model. However, we would emphasise that the confidence bias we simulate here is likely distinct to that induced by positive evidence. More specifically, such changes in metacognitive bias here occur due to shifts in parameters governing higher-order nodes of the circuit (UM). We think that these two instances of confidence modulation in absence of d’ change may index different metacognitive “biases” in the system – at first-order and higher-order levels, respectively.

**Supplementary Note 6: Experiment 1 fit with holdout**

To demonstrate that our model fits generalise to predict unseen test data, we re-ran our fitting algorithm on all Experiment 1 participants – holding out a 30% random non-stratified partition of each participant’s data for validation, and fitting on the remaining 70%.

Supplementary Figure 5 below shows that the model generalises well in the case of choice accuracy (A) and response times (B).
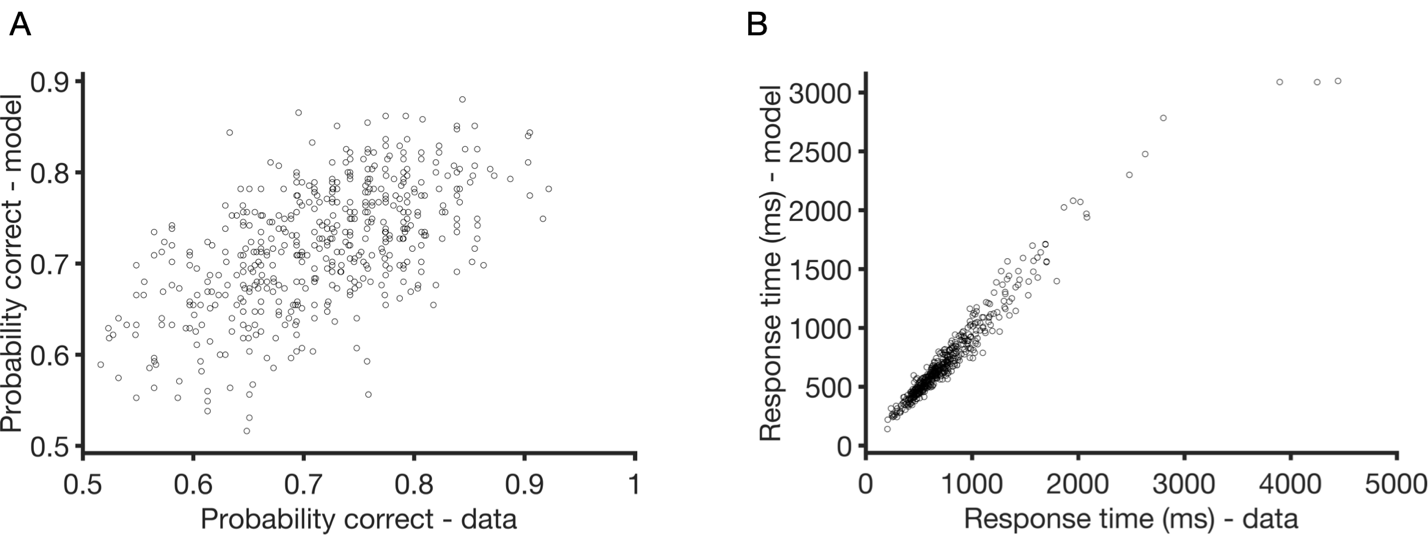

**Supplementary Figure 5.**  **Model parameters fitted to a subset (70%) of the data were then used to simulate data for each participant.** Empirical overall accuracy and mean response time were calculated from the holdout (30%) set unseen by the model during fitting**.** The model fits participants’ overall accuracy (A) and response time (B).

**Supplementary Note 7: UM as a low dimensional marker of metacognitive profile**

We have run a regression where mean empirical confidence for Experiments 1 and 2 was entered as a predictor of fitted UM, controlling for gender, IQ and age. For experiment 2, mean empirical confidence was indeed positively correlated with UM, despite the distribution matching approach (Beta = 0.12, SEM = 0.04, p<0.005). A similar positive correlation between UM and mean empirical confidence was observed in Experiment 1 (Beta = 0.0017, SEM = 0.04), but this did not reach significance (p>0.5). Overall, these findings suggest that such fitted parameters may provide insight into a subjects’ metacognitive profile even in the absence of confidence rating data.

**Other Supplementary Figures:**

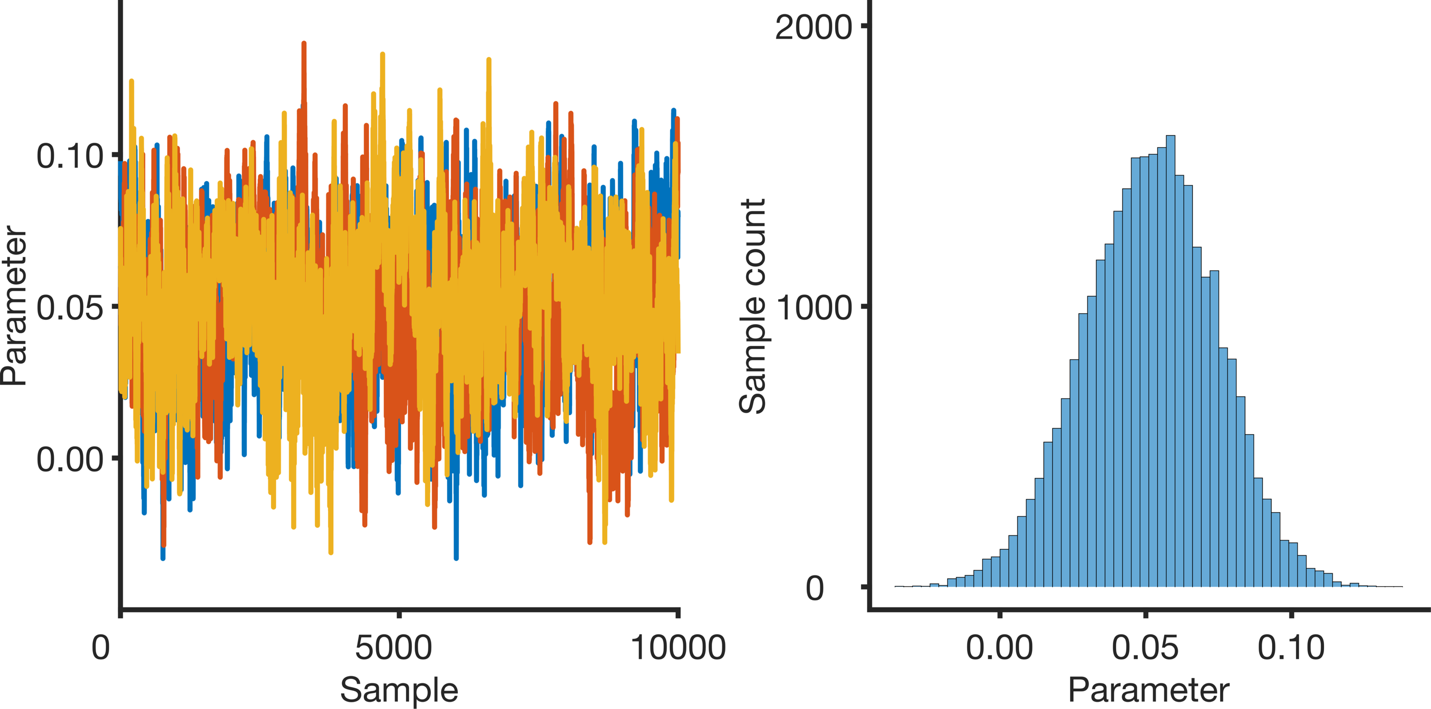

**Supplementary Figure 6. Hierarchical estimation of the impact of fitted UM on observed meta_d’/d’ ratio** **using a simultaneous regression approach with the UM parameter as a covariate (Harrison et al., 2020).** A positive association between the strength of uncertainty modulation and metacognitive efficiency (mean value of mu_beta = 0.0516, 95% highest density interval = (0.0813, 0.0016)).

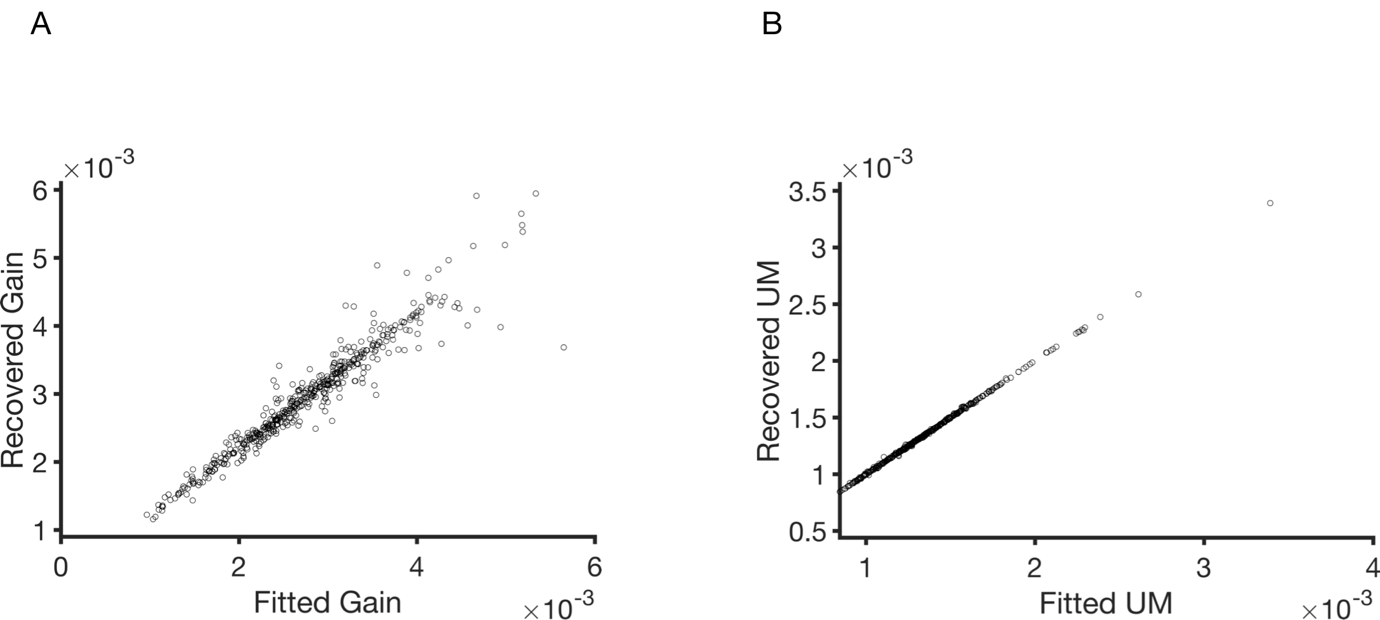

**Supplementary Figure 7.** **Results from parameter recovery simulations for experiment 1 for both (A) Gain and (B) UM**. Recovery simulations were done by simulating the fitted parameter for each participant, and then fitting to synthetically generated data (i.e. accuracy and mean response time). Both figures highlight a reasonably stable fit.

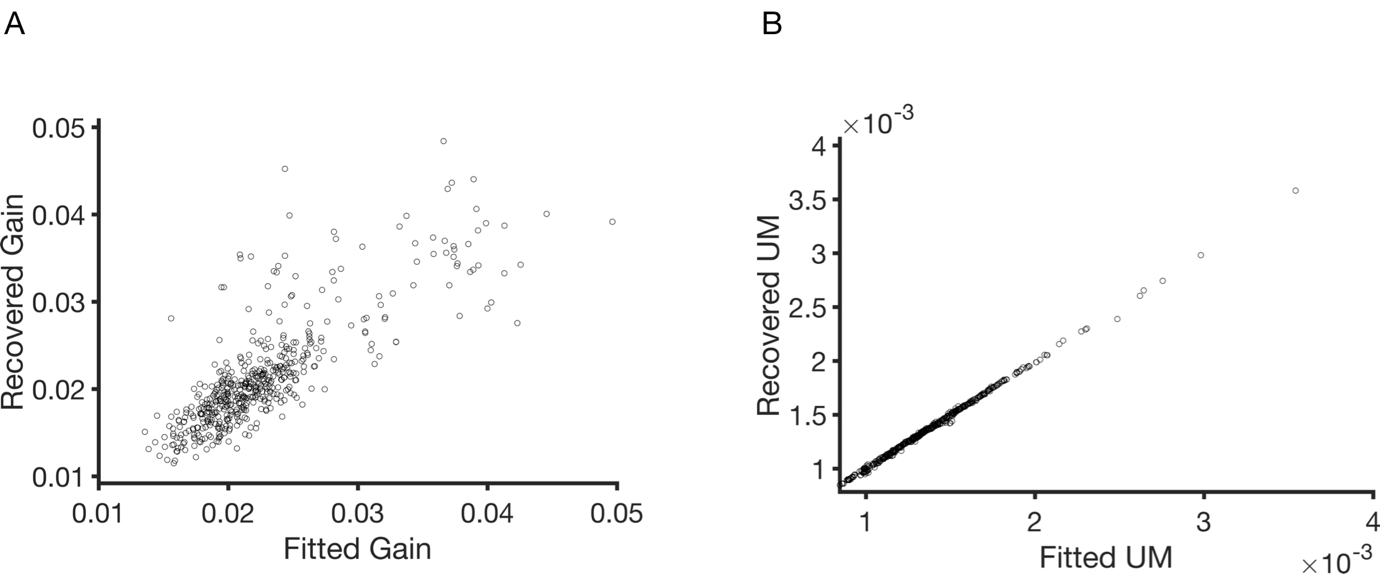

**Supplementary Figure 8.** **Results from parameter recovery simulations for experiment 2 for both (A) Gain and (B) UM**. Gain recovery for experiment 2 fits is more challenging, which could be partly due to the lower variance in difficulty experienced by each participant in experiment 2 (See Supplementary Figure 7).

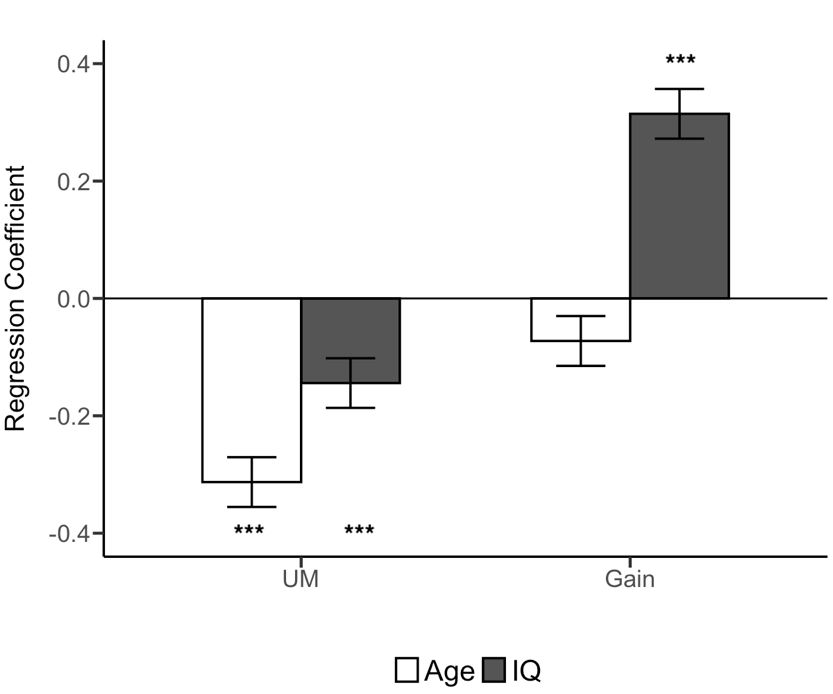

**Supplementary Figure 9.** **Relationship between Age/IQ and model parameters.** Age and IQ data from experiment 1 were entered into a multiple regression model (controlling for gender) predicting the strength of uncertainty modulation and gain parameters from the model fits to task performance (choices and response times). The results reveal that higher IQ is significantly associated with increased gain – suggesting a strong relationship between IQ and accuracy. The relationship is between IQ and Uncertainty Modulation (UM) is weaker, but still significant. Older age is associated with weaker UM.

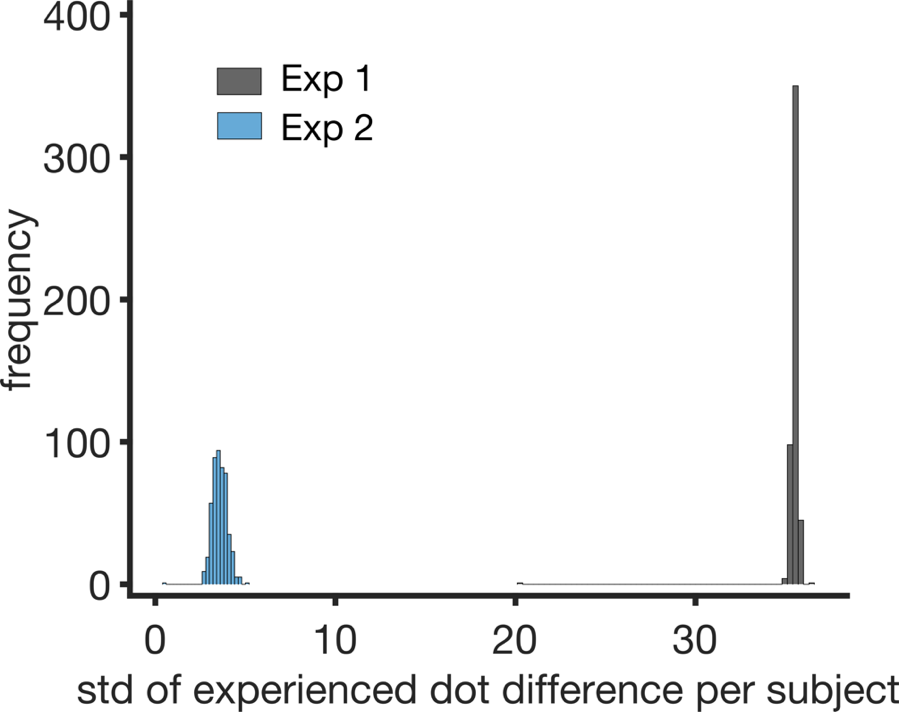

**Supplementary Figure 10. Variance in difficulty experienced by participants in Experiment 1 and 2.** Distribution of standard deviation of experienced dot difference per subject for experiment 1 (black) and experiment 2 (blue). Variance in difficulty between participants is much lower in experiment 2 compared to experiment 1 due to the use of a staircase procedure in the former.

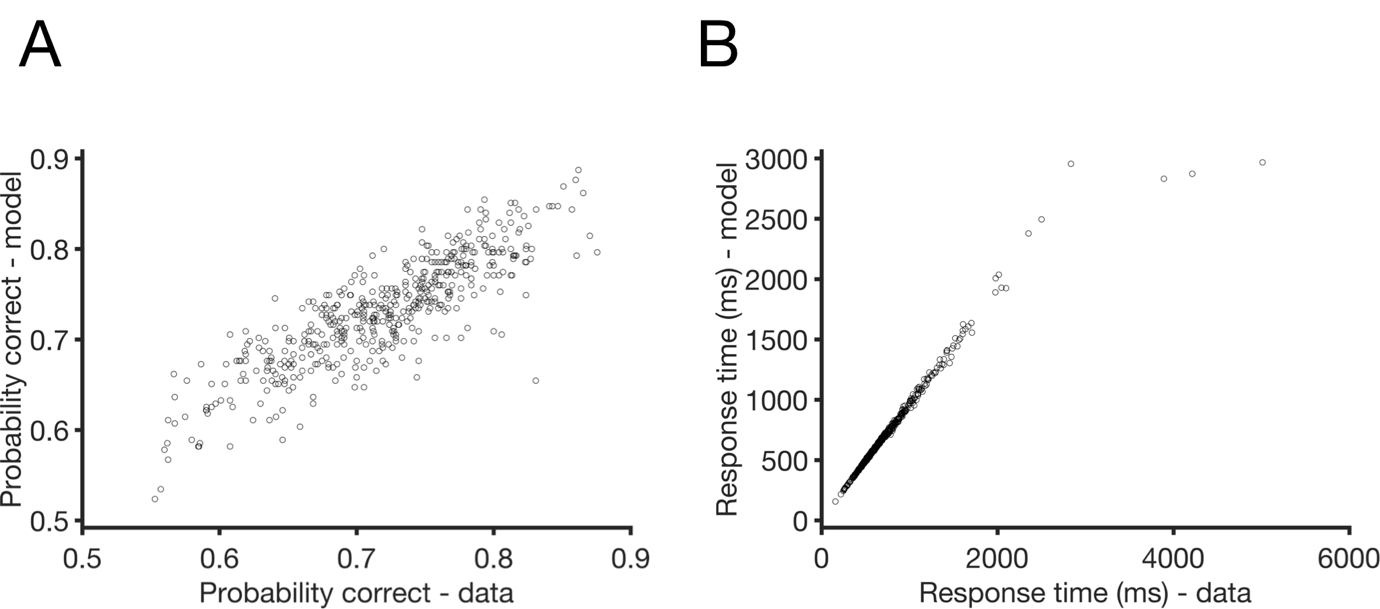

**Supplementary Figure 11. (A) Individual accuracy and (B) mean response time model fits for Experiment 1 without resetting the random number generator seed.**

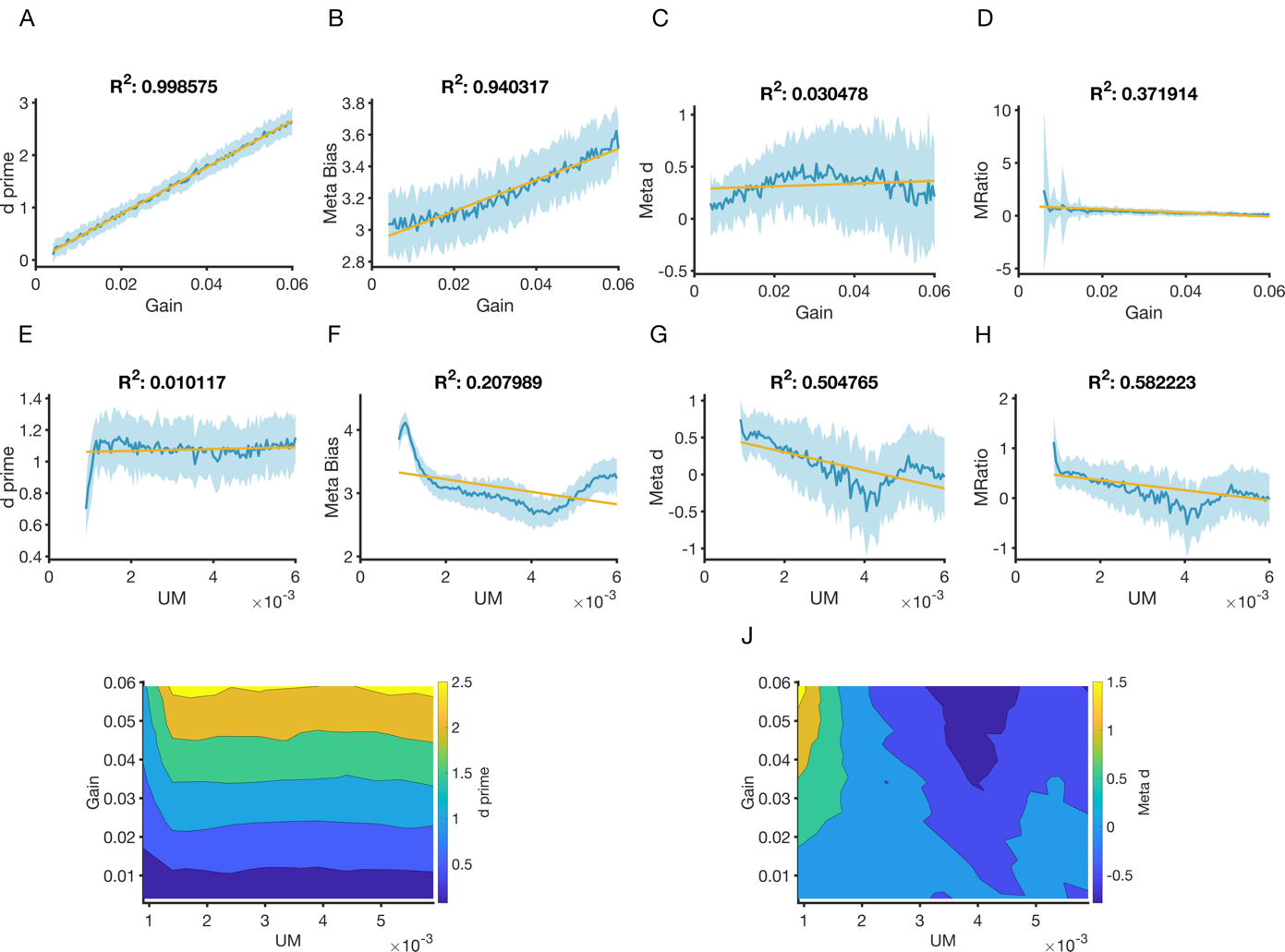

**Supplementary Figure 12.**  **Simulations of Figure 2 in the main manuscript with alternative parameter values.** In simulations (**A-H**), where the gain (UM) parameter is varied, UM (gain) was fixed at 0.0015 (0.0029). In this parameter subspace, meta_d’/d’ ratio is low – particularly in relation to d’.
